# Quantitative insights into age-associated DNA-repair inefficiency in single cells

**DOI:** 10.1101/628909

**Authors:** Thomas Z. Young, Ping Liu, Murat Acar

## Abstract

The double strand break (DSB) is a highly toxic form of DNA damage that is thought to be both a driver and consequence of age-related dysfunction. Although DSB repair is essential for a cell’s survival, little is known about how DSB repair mechanisms are affected by cellular age. Here we characterize the impact of cellular aging on the efficiency of single-strand annealing (SSA), a repair mechanism for DSBs occurring between direct repeats. Using a single-cell reporter of SSA repair, we measure SSA repair efficiency in young and old cells, and report a 23.4% decline in repair efficiency. This decline is not due to increased usage of non-homologous end joining (NHEJ). Instead, we identify increased G1-phase duration in old cells as a factor responsible for the decreased SSA repair efficiency. We further explore how SSA repair efficiency is affected by sequence heterology and find that heteroduplex rejection remains high in old cells. Our work provides novel quantitative insights into the links between cellular aging and DSB repair efficiency at single-cell resolution in replicatively aging cells.

## INTRODUCTION

DNA damage has long been hypothesized to be both a driver and consequence of aging^1, 2^. Old tissues and cells accumulate DNA damage and mutations^3, 4, 5^. Such mutations can negatively impact tissue homeostasis and organismal function, and may even increase the risk of cancer^6, 7, 8^. The DNA double strand break (DSB) represents a major category of DNA damage. Inability to repair a DSB can lead to cell death or genomic instability^9^. Even when a DSB is repaired, there are mutagenic repair mechanisms that can both increase genetic variability and impact cellular fitness^10^. Whether cellular aging impairs DSB repair efficiency in single cells remains unknown.

A useful model system to understand how cellular aging affects DSB repair efficiency is that of replicative aging of the budding yeast *Saccharomyces cerevisiae*^11^. Replicative lifespan is the number of times a yeast mother cell produces daughters^11^. This number varies across genetic backgrounds and growth conditions, and is connected to several metabolic pathways^12, 13^. The connection between DSB repair and replicative lifespan is important for several reasons. To avoid chromosomal instability, most cells that cannot repair a DSB will halt division, and thus have a limited replicative lifespan^14^. In the other direction, continued cell division after mutagenic DSB repair results in propagation of these mutations to future generations with potentially negative consequences on cell fitness.

Multiple mechanisms exist to repair double strand breaks and their efficiency and ability to function properly could change with replicative age. Among these mechanisms, the non-homologous end joining (NHEJ) pathway involves ligation of the free ends flanking the double strand break^15^. The other main class of DSB repair mechanisms are the homology-directed repair (HDR) mechanisms, which make use of sequence homology between the break-site and a repair template. Regulation of homology-directed repair can have significant consequences for the genome. This is due to the ability of different HDR mechanisms to change allele copy numbers and lead to recombination between chromosomes. The relative usage and efficiency of different DSB repair mechanisms depend on factors including cell-cycle stage, ploidy, and cell type^15, 16, 17, 18^. Previous studies have detected changes in repair pathway usage between chronologically old and young tissues, that could indicate age-related changes in repair efficiency^19, 20^. However, how the repair efficiency of specific DSB repair pathways longitudinally changes in mitotically aging single cells remains unexplored.

Here we assess whether the efficiency of DSB repair via the single-strand annealing (SSA) pathway changes with the age of the host cell. The SSA pathway repairs double strand breaks occurring between direct repeats of an identical sequence, resulting in deletion of the intermediate sequence (Fig. 1a)^15^. Repetitive sequences play important roles in cellular function, with the rDNA locus being one prominent example^21, 22^. An age-related change in the efficiency of SSA, which would lead to differences in the copy numbers of the repeated sequence, would be expected to have important consequences for the cell. Since SSA and other homology-directed repair pathways share regulatory aspects such as end-resection and Rad52 recruitment to the repair sites, understanding how SSA efficiency changes with age can provide key insight into whether the efficiency of other homology-directed repair mechanisms would also change with age. Using a singlecell longitudinal approach^23^ in a haploid genetic background, we measure the efficiency of single-strand annealing repair in young and older cells. We further explore age-related changes in SSA repair efficiency as they relate to age-related changes in cell cycle, non-homologous end joining pathway activity, and in terms of the amount of heterology between the SSA repeats.

**Figure 1.**
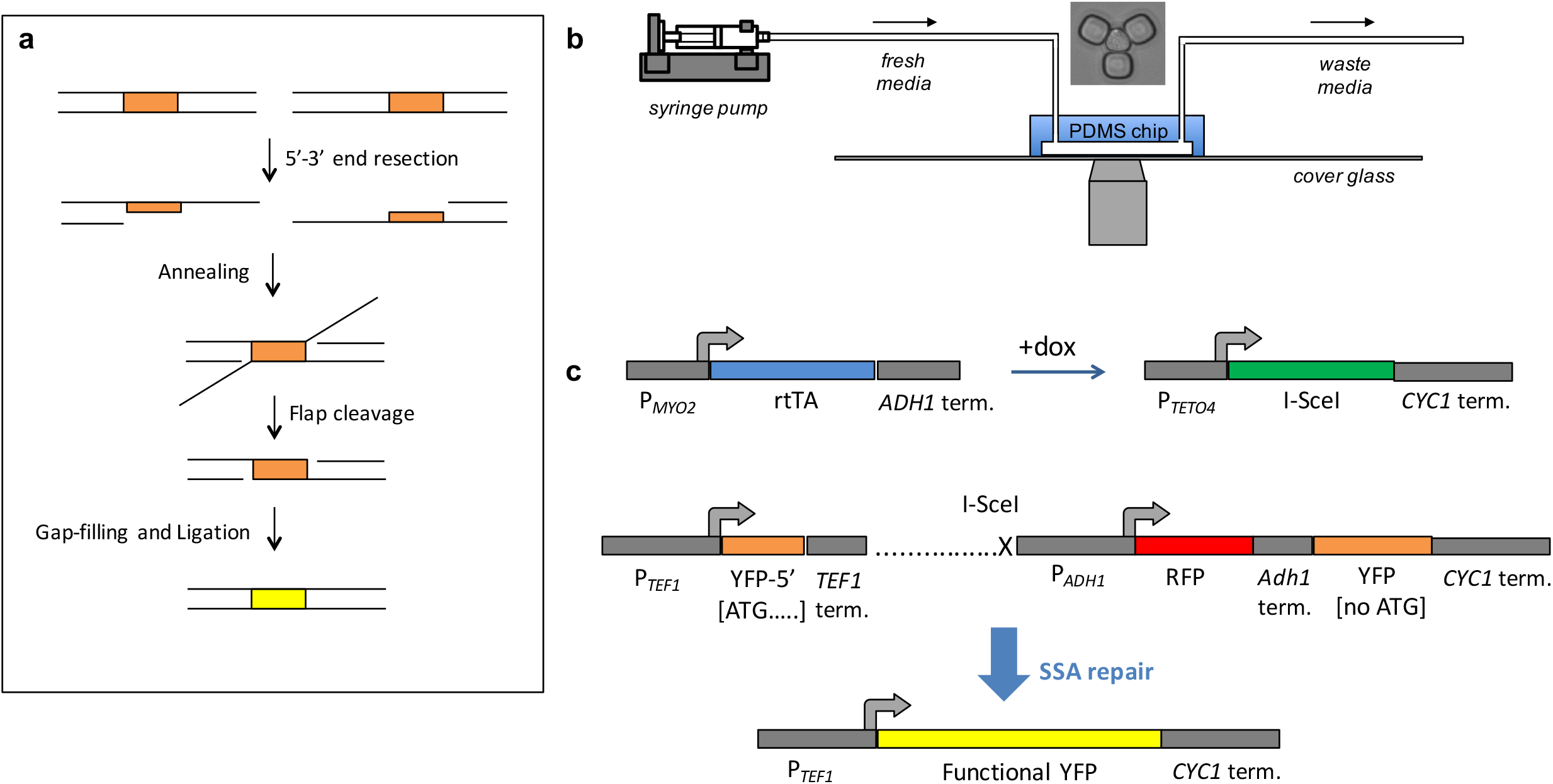
Schematics of the experimental system and SSA repair measurements. **a.** Diagram showing the steps of single-strand annealing based repair. The direct repeats are highlighted in blue. The upper row shows the situation immediately after the double strand break. **b.** Schematic showing the setup of the experiment. The yeast replicator device allows replicative aging of yeast to be observed using a microscope. Fresh media is supplied, and waste removed over the course of the movie. A microscope image of a trapped yeast cell is shown above the chip. **c.** Genetic constructs used to measure SSA. rtTA is expressed from the constitutive P*_MYO2_* promoter. In the presence of doxycycline, rtTA activates expression of I-SceI from a non-leaky P*_TETO4_* promoter. I-SceI cutsite consists of a pair of inverted 18 bp I-SceI sites placed within the SSA reporter. The non-fluorescent 5’ YFP repeat contains only the 5’ 192 bp of YFP. The 3’ YFP repeat is not expressed because it consists of the entire YFP ORF except for the start codon. Between the *TEF1* terminator and 3’ YFP repeat is a 2.1 kb stretch of DNA (dotted line), with inverted I-SceI cut sites at a distance of 0.5 kb from the *TEF1* terminator. Between the I-SceI cutsite and the 3’ non-functional YFP is an *ADH1* promoter driving mCherry (abbreviated by ‘RFP’). Degron-tagged RFP expression below a threshold was used as a reporter for DNA cutting. After SSA repair occurs between the repeats, the resulting product is a full length YFP with a start codon.

## RESULTS

### Developing a single-cell SSA repair reporter and experimental design

Cells were aged in a microfluidic chip (Fig. 1b) to obtain either young or old cells in which to measure SSA efficiency. To control the age at which SSA repair is assessed, we used a haploid strain in which the endonuclease I-SceI is expressed under the doxycycline-inducible P*_TETO4_* promoter. To measure SSA, we developed an SSA reporter containing an I-SceI cut site between two nonfunctional YFP repeats integrated at the chromosomal ura3 locus. If SSA repair occurs using the two YFP repeats of the reporter, then a functional YFP is formed and detected in single cells aging in real-time (Fig. 1c). For direct detection of whether cutting occurred or not, an RFP reporter is constitutively expressed from the genomic region between the two non-functional YFP repeats; cutting is detected by the dilution of RFP concentration that results from a halt in RFP expression during continued cell growth. The I-SceI cut-site consists of two adjacent 18 bp I-SceI recognition sequences in inverted orientation so that after two I-SceI cleavages the DNA ends flanking the break are incompatable^24^. This situation is more representative of naturally occurring DSBs^24^. We also used a control strain identical to the strain carrying the SSA repair cassette, except for the absence of the I-SceI cut site.

For inducing I-SceI expression and cutting, we chose a 4-hour time window during which cells were exposed to doxycycline (dox). SSA repair efficiency was measured in separate experiments targeting young or older cells. Repair efficiency was quantified using the fraction of initially YFP^-^ cells that became YFP^+^ within the 9 hour period (4 hours of dox treatment plus 5 hours of post-dox inspection) after addition of dox. A cutoff-based approach was used to determine whether a cell was YFP^-^ or YFP^+^ at any given time (Fig. S2). Only cells that were observed to be alive within the 5-hour post-dox time window were considered. For the SSA repair experiments in young cells, the average age of those cells was 2.6 generations at the beginning of the dox treatment. In the experiments with old cells, the average age was 18.1 generations at the beginning of the dox treatment (Fig. 2a, Fig. S1), which is close to the average age of the haploid yeast at ~23 generations^25^. The two expected phenotypes of YFP appearance, or its absence, were observed and quantified across individual cells longitudinally (Fig. 2b, Fig. S3a,b, Fig. S4).

**Figure 2.**
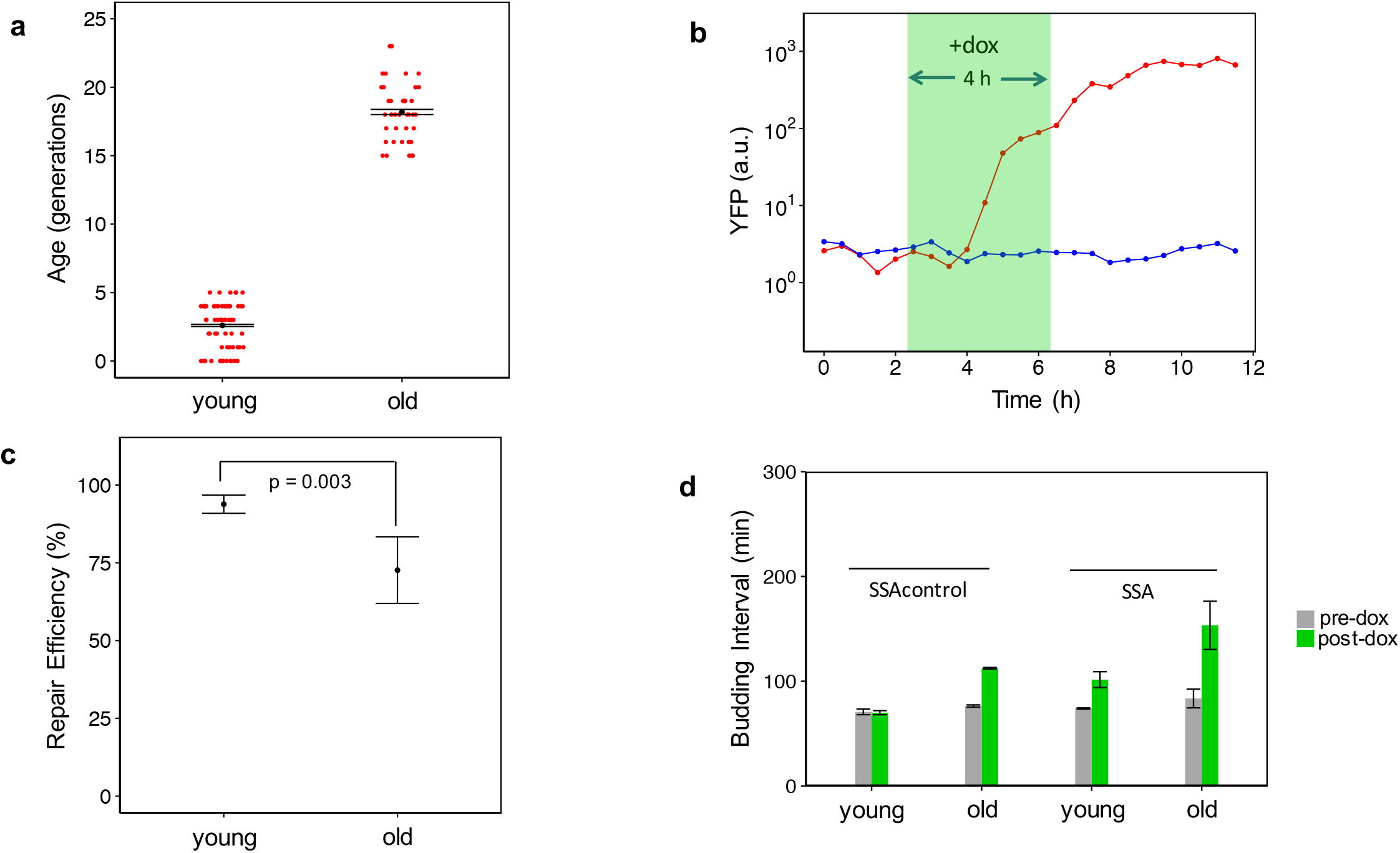
Measuring SSA repair efficiency under perfect sequence homology. **a.** Points represent the age of each cell at the beginning of the doxycycline treatment. The mean age of old cells (N=2 replicates) was 18.1 generations, while the mean age of the young cells (N=2 replicates) was 2.6 generations. **b.** YFP trajectories are shown for two young sample cells, with the cell with the red trajectory representing successful SSA repair. **c.** The fraction of cells with successful SSA repair was calculated for each experimental replicate. Shown are mean+/-SEM (N=2) by age group for the fraction of repaired cells. The old cell replicates had a mean repair efficiency of 71.8% compared to a mean repair efficiency of 93.8% for the young cell replicates. The p-value of 0.003 is for a twosided Fisher’s Exact Test for association between repair and age on pooled data. **d.** Shown are the mean+/-SEM (N=2) of the average budding intervals measured from individual cells of the two replicates before and after doxycycline addition. Pre-dox corresponds to the 4-hour time window before doxycycline addition. Post-dox corresponds to the 9-hour time window after doxycycline addition. At both young and old ages, the strain carrying the SSA repair reporter exhibits increased budding interval relative to the control strain, indicative of DNA damage-induced cell-cycle arrest (from 74+/-1 min to 102+/-8 min in young cells with the cutsite; from 84+/-9 min to 153+/-23 min in old cells with the cutsite).

### Measuring SSA repair efficiency in young and old cells

The SSA repair efficiency for the old cells was significantly lower than that of the young cells (71.8% vs. 93.8%), indicating an age-associated decline in SSA repair (Fig. 2c; *P*-value 0.003, Fisher’s exact test). The fraction of unrepaired old cells that were repaired in the subsequent 5-hour time window after dox removal was also found to be low (1/11) (Fig. S5). Therefore, the absence of repair at the 9-hour time point was not simply due to a small reduction in the speed of repair. Additionally, the average time after dox addition at which YFP first appeared was similar between the old and young repaired cells, indicating that aging does not significantly affect the speed of SSA repair in cells that are able to carry out SSA repair (Fig. S4). To assess whether cells were arresting their cell cycle due to cutting by I-SceI, we calculated the average budding time experienced by cells in the 4-hour time interval before dox addition, and the 9-hour time interval after dox addition. We compared this to the corresponding average budding times calculated for the control strain lacking an I-SceI cut site. In the time window prior to doxycycline treatment, the two strains show similar average budding times (Fig. 2d) for the same age groups. Consistent with I-SceI cleavage resulting in cell-cycle arrest, average budding times after dox addition are higher in the strain containing the cutsite, compared to the control strain without the cutsite. After dox addition, the strain containing the cut-site had an average budding time of 153±23 min in old cells compared to 112 ± 1min for the old cells of the control strain (Fig. 2d). Therefore, induction of the DSB leads to longer cell-cycle durations on average.

To determine whether the unrepaired YFP^-^ cells were arrested in their cell cycle consistent with cutting and failure to repair, we considered the time that had passed since these cells last started forming a daughter bud. Since the first appearance of a daughter bud corresponds to early S-phase, the amount of time since the previous bud initiation can include the G1 phase of the subsequent cell cycle. If the cells are arrested in G2/M phase, on the other hand, this time period only includes the S/G2/M phase of the same cell cycle. Compared to the YFP^+^ repaired cells, cells that were YFP^-^ at the 9^th^-hour time point after doxycycline addition had not budded for a longer period of time (Fig. S6). These unrepaired cells are therefore either arrested for a long time in G2/M or went through mitosis before experiencing a long G1 in the subsequent cell cycle.

To determine whether the cells that failed to produce YFP by SSA repair exhibited any cell cycle delays consistent with a DSB-induced cell cycle arrest, we measured the average budding interval of each cell in three non-overlapping time windows: the 4-hour window prior to doxycycline treatment, the 4-hour window coinciding with doxycycline treatment, and the 5-hour window after doxycycline treatment. The old YFP^-^ cells exhibited slightly longer average budding intervals prior to doxycycline addition compared to the old cells that were eventually repaired (Fig. S7a). During the 4-hours of doxycycline exposure and the 5-hour period after doxycycline exposure, both YFP^+^ and YFP^-^ cells showed an increased average budding interval compared to the control strain (Fig. S7b, S7c). Average budding intervals in the 5-hour post-doxycycline time window for young YFP^-^ cells were far longer than those for young cells of the SSA control strain lacking a cutsite (Fig. S7c). These results suggest that cutting efficiency is 100% efficient in young cells. For old cells, on the other hand, the considerable overlap in average budding interval for YFP^-^ cells and control cells lacking a cutsite makes budding interval a poor measure of cutting efficiency (Fig. S7c), necessitating the use of more direct approaches – such as the RFP reporter used in this study – for determining cutting efficiency.

### Direct measurement of I-SceI cutting efficiency in old cells

The age-associated decline in SSA repair efficiency could be due to several factors. First, the cells could be cut and failing to complete repair using SSA. Second, weaker induction of I-SceI in old cells could result in a lower rate of cutting. A third possibility is that the cells in which YFP fails to appear are actually cut by I-SceI, but repaired by a different repair pathway like NHEJ or MMEJ (Microhomology Mediated End Joining) that does not generate functional YFP. To address the possibility of decreased cutting efficiency by I-SceI in old cells, we constructed a strain in which the RFP originally introduced between the two non-functional YFP repeats was further tagged with a degron. The degron tag we used is a truncation of the C-terminal destabilization motif from the cyclin protein Cln2 and it leads to the fast degradation of Cln2 and RFP. The stability of the original non-tagged RFP results in poor detection of any cutting due to the longer time and continued growth requirement for the detection of RFP loss after cutting. In the strain carrying the degron-tagged RFP, a halt in RFP transcription due to cutting by I-SceI is expected to result in a rapid and prolonged decline in RFP transcription. At the 9^th^-hour time point after doxycycline addition, we measured the length of time that each cell had been RFP^-^ using a fixed cutoff for RFP (Fig. 3a, Fig. S8). Soon after addition of doxycycline, young and old cells of this strain showed a rapid and lasting decline in RFP expression that did not occur in old cells of a control strain containing the RFP-degron fusion, but without a cutsite (Fig. 3b, Fig. S3h,i,j). Since old cells of the cuttable and control strains, both containing the SSA cassette and degron-tagged RFP, have similar RFP expression levels prior to the decline in RFP expression, we conclude that the difference in the metric of “RFP^-^ duration” is not due to differences in promoter activity (Fig. 3c). Only one of 58 (1.7%) old cells of the cutsite-free strain containing the RFP degron lost RFP expression for over 4.5 hours. 98% of all old, initially YFP^-^ cells from the SSA strain carrying degron-tagged RFP showed either a decline in RFP signal to background levels for over 4.5 hours, or production of functional YFP. Based on these results, we concluded that I-SceI cutting in old cells was very efficient. Therefore, the observed decline in SSA repair efficiency in old cells compared to young cells is due to either decreased SSA repair efficiency or increased efficiency of competing repair pathways.

**Figure 3.**
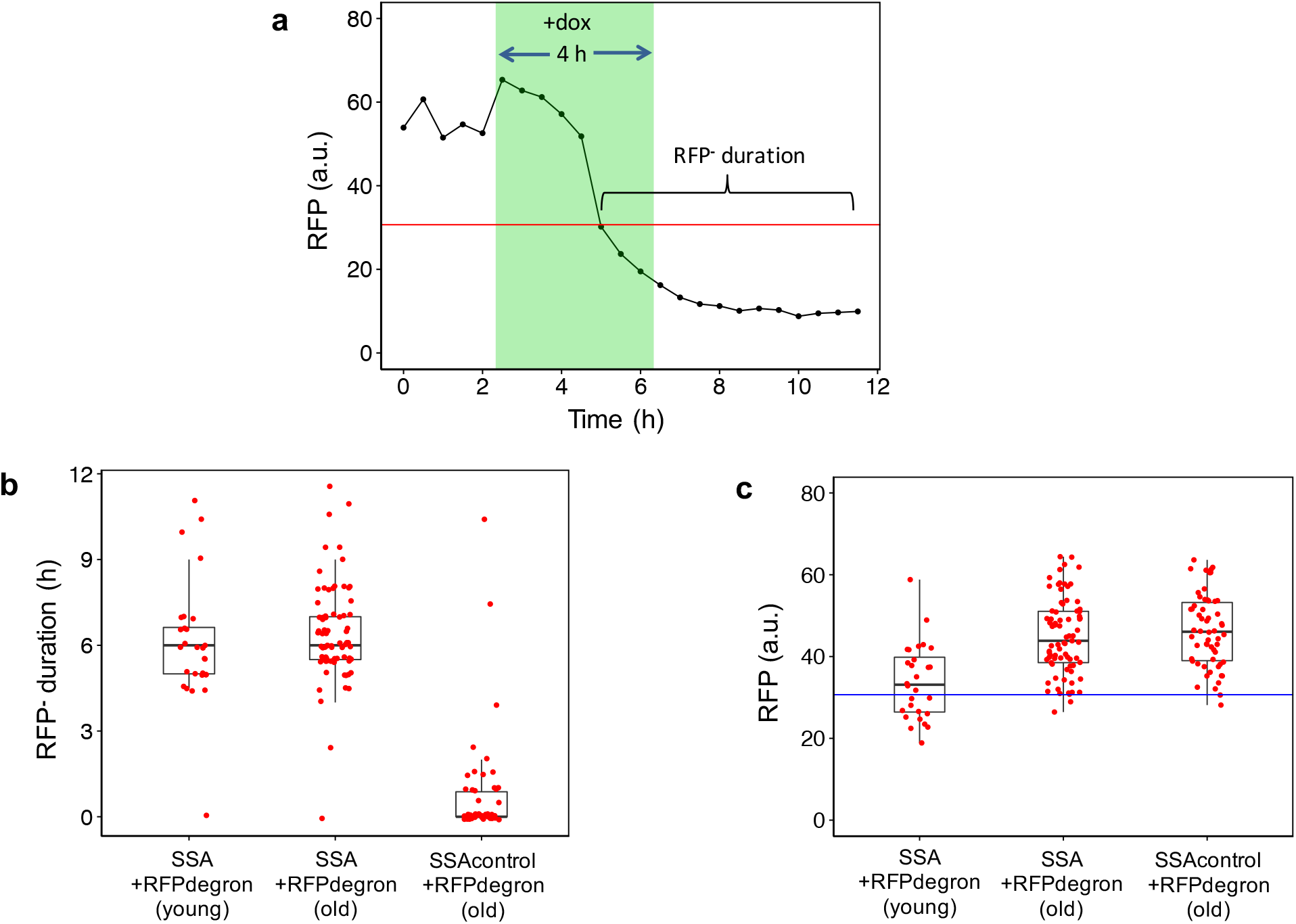
Directly measuring I-SceI cutting efficiency using a degron-tagged mCherry (‘RFP’) in the SSA repair cassette. **a.** An RFP cutoff was applied to cells from two strains containing a degron-tagged mCherry (abbreviated ‘RFP’) within an SSA repair reporter: one strain contained an I-SceI cutsite while the other lacked the cutsite. For each cell analyzed, the duration of RFP absence was quantified between the first occurrence of below-cutoff RFP level and the end of the 9-hour post-doxycycline time window. **b.** The duration of RFP absence is shown for old and young cells of the strain containing a degron-tagged RFP within an SSA repair reporter and old cells of the same strain missing the I-SceI cutsite. 43 of the 58 old cells (corresponding to 74%) analyzed from the second strain missing the cutsite had a duration of RFP absence quantified as 0 min, compared to 1 out of the 79 old cells from the first strain carrying the cutsite. Using pooled data from two independent experiments, the box plots show the distribution of the single-cell RFP^-^ durations for each cell. **c.** For each cell that was missing RFP (expression below the cutoff) at the 9^th^ hour time point after doxycycline addition, the RFP values of each cell measured prior to their being below cutoff were measured and averaged for each cell. Using pooled data from two independent experiments, the box plots show the distribution of the single-cell-averaged RFP values corresponding to each cell’s RFP^+^ period. The RFP values corresponding to the RFP^+^ period are similar between old cells of the two degron-tagged SSA reporter strains, with and without the cutsite.

### Measuring age-associated SSA repair efficiency in the absence of NHEJ

We hypothesized that the age-related repair deficiency in the SSA repair strain was due to increased competition from the non-homologous end joining repair pathway in old cells. Previous work performed in young cells has shown that DSB repair is more biased towards NHEJ during G1 phase when there is no identical sister chromatid for homology-based repair^26^. Deletion of the key NHEJ proteins Ku and Dnl4 has also been shown to increase DSB end-resection, an important step of homologous recombination based repair pathways^27^.

If the decline we observed in the fraction of YFP^+^ cells was due to NHEJ outcompeting SSA in old cells, and SSA itself is still fast and efficient, then inhibition of NHEJ would increase the SSA repair efficiency in old cells back to that of young cells. To test for this possibility, we generated a perfect-homology SSA repair strain identical to the previous strain except with the non-homologous end-joining gene *DNL4* deleted. The Dnl4 protein is a DNA ligase that is an essential component of the NHEJ mechanism. Deleting *DNL4* does not affect replicative lifespan^28^. Using this strain, we performed the same type of doxycycline-induction experiments by targeting young (average age of 2.4 generations) or older cells (average age of 19.3 generations) (Fig S1), and measured the fraction of cells with YFP fluorescence as a sign of SSA repair (Fig. S3e, S3f). SSA repair efficiency in young *dnl4Δ* cells was 90.4% (Fig. 4a), essentially unaffected by the deletion compared to the non-deleted strain’s 93% repair efficiency. In old *dnl4Δ* cells, we measured a 65.0% SSA repair efficiency compared to the 70.0% SSA repair efficiency in the old cells of the non-deleted strain (Fig. 4a). As with the *DNL4* intact strain, only a small fraction of the *dnl4Δ* cells that were unrepaired at the 9-hour post-doxycycline time point were repaired in the subsequent 5 hours (Fig. S5). In the 9 hours after doxycycline addition, the average budding time was 156±16 min for old cells and 102±18 min for young cells, similar to the corresponding values for the strain with intact *DNL4* (Fig. 4b). In this strain, the ratio of the repair efficiency in old cells to the one in young cells was 0.719, very similar to the ratio measured in the SSA strain with intact *DNL4* (Fig. S11). Consistent with this, the ratio of average budding interval in the 9-hour post-doxycycline time window between old and young cells of this strain was also similar to the ratio for the strain with intact *DNL4* (Fig. S12). The lack of increase in the SSA repair efficiency when the NHEJ repair mechanism is inactive suggests that increased usage of NHEJ is not the cause of the age-specific inefficiency in SSA-based repair.

**Figure 4.**
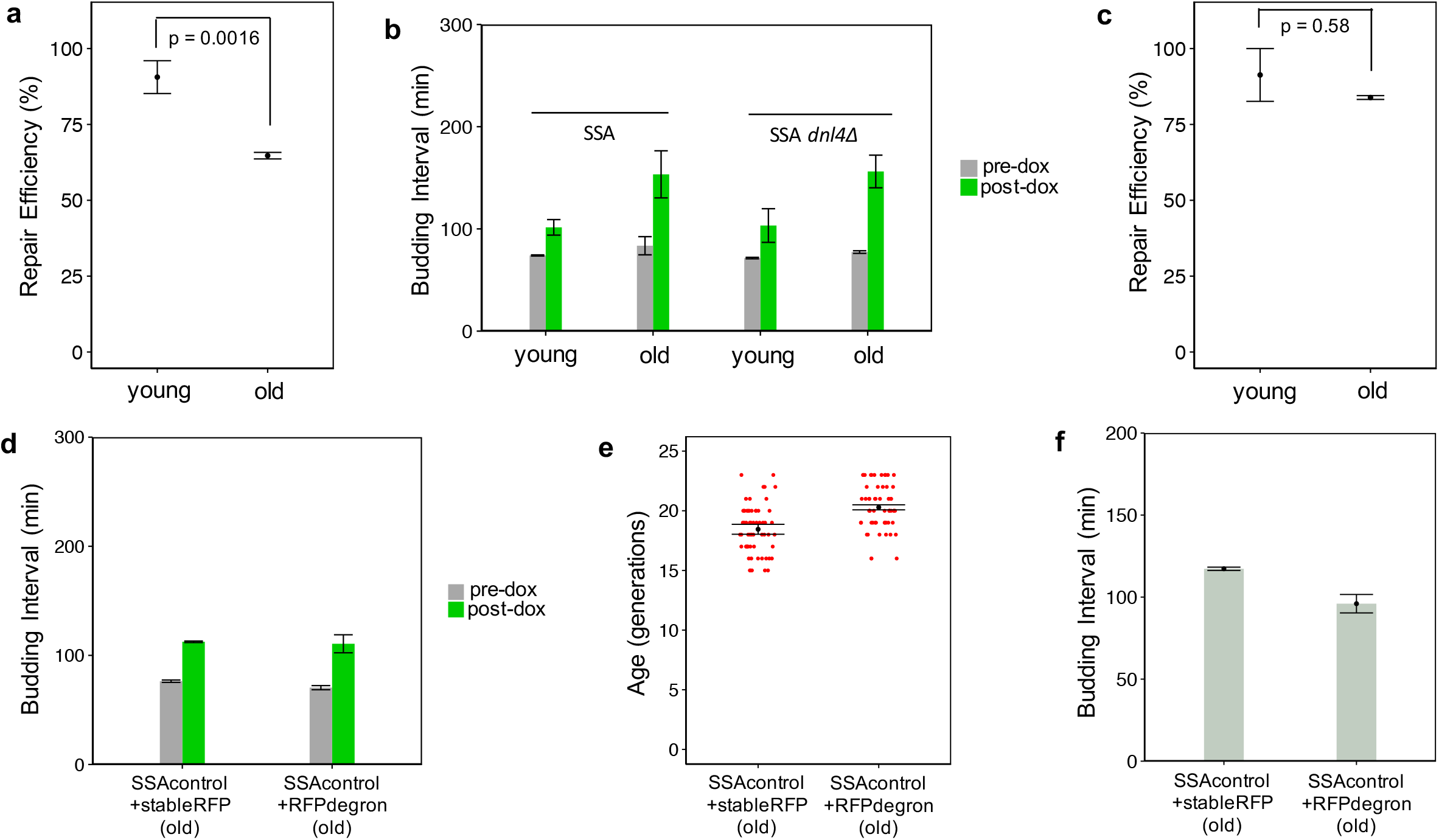
The age-associated decline in SSA repair efficiency is due to changes in cell-cycle progression in old cells. **a.** *dnl4Δ* cells fail to show an increase in SSA repair efficiency when old. The mean repair efficiency for old cells (N=2 replicates) was 65.0%, while for young replicates (N=2), it was 90.4%. The mean and SEM (N=2) of these replicate-based statistics are shown. The p-value of 0.0016 is for a two-sided Fisher’s Exact Test to test for association between repair and age. **b.** Shown are the mean+/-SEM (N=2) of the average budding intervals measured from individual cells of the two replicates during the 4-hour pre-doxycycline addition, and 9-hour post-doxycycline addition time window. The *dnl4*Δ strain shows a large increase in budding interval after doxcycycline treatment regardless of age (from 71+/-1 min to 103+/-16 min in young cells; from 77+/-1 min to 156+/-16 min in old cells). **c.** The strain containing a degron-tagged mCherry (‘RFP’) within the SSA repair reporter has higher SSA repair efficiency in old cells. **d.** Average budding intervals (‘times’) during the 9-hour post-doxycycline time window are similar in old cells of the SSAcontrol strain (no cutsite) carrying degron-tagged mCherry (‘SSAcontrol+RFPdegron’), and old cells of the SSAcontrol strain carrying stable mCherry (‘SSAcontrol’). Shown are mean+/-SEM (N=2) of the average budding intervals measured from individual cells of two replicate experiments, pre-dox (during the 4-hour time window before doxycycline addition) and post-dox (during the 9-hour time window after doxycycline addition). In old cells, the average post-dox budding interval was 115+/-6 min for the RFPdegron-carrying control strain, and 111+/-8 min for the stable-RFP-carrying control strain. **e.** Single-cell ages at the start of doxycycline treatment for the old age groups of the SSAcontrol strains (no cutsite) with stable RFP or degron-tagged RFP (for which average budding intervals were calculated in this panel). Mean+/-SEM (N=2) of replicate-based averages are overlaid on single-cell age values pooled from two replicates; the average age is 18.5 generations for the strain with stable RFP, and 20.3 generations for the strain with the degron-tagged RFP. **f.** Average budding intervals measured for old cells of the SSAcontrol strains (no cutsite), with stable RFP or degron-tagged RFP. Only generations larger than 15 were considered for cells of age 15 generations or older at the time of doxycycline addition. Mean+/-SEM (N=2) of replicate-based averages of single-cell-averaged budding intervals are shown. The mean values are 117+/-1 min for the strain with the stable RFP, and 96+/-6 min for the strain with the degron-tagged RFP.

### The role of changes in cell-cycle progression on SSA repair efficiency in young vs old cells

Since eliminating Dnl4 did not restore SSA repair efficiency in old cells to the levels observed in young cells, we next considered whether some aspect of SSA repair itself was deficient in old cells. SSA falls under the class of homology-directed repair mechanisms that are normally inactive in the G1 phase, and activated during other phases of the cell cycle. It has been shown that several generations prior to the end of their replicative lifespan, cells undergo a lengthening of the cell cycle^29, 30, 31^. This increase in cell cycle duration correlates at the single-cell level with an increase in the duration of G1 phase, and lower expression of key G1/S transition proteins including the cyclin Cln2^30^. Outside of G1 phase, the cyclin dependent kinase subunit Cdc28 is activated by S- and G2-phase cyclins (Clbs) to promote homology-directed repair. Clb-activated Cdc28 promotes HDR by inhibiting the binding of the NHEJ-promoting Ku proteins to the DSB break site and by promoting end-resection and recruitment of HR proteins to the site^10, 26, 27^. An age-related increase in G1 phase duration could result in a decrease in SSA repair efficiency due to reduced levels of Clbs that activate Cdc28’s HDR-related activities^10^. Since the Ku complex prevents end-resection in the absence of Cdc28 activity, deletion of *DNL4* under this scenario would be expected to fail to activate homology-directed repair by SSA, which was observed (Fig. 4a). Consistent with this hypothesis, we saw that old cells containing the SSA cassette that failed to repair YFP had longer average budding intervals in the 4-hour time window prior to doxycycline treatment compared to old cells that were able to repair YFP (Fig. S7a, S9).

Interestingly, we observed that SSA repair efficiency in old cells of the SSA strain containing the degron-tagged RFP was 83.6%, a number higher than the efficiency observed in the old cells of the SSA strain containing the stable RFP without the degron (Fig. 4c). Additionally, the speed of SSA repair in old cells of this degron-carrying strain was fast – YFP was detected sooner in old cells of this strain than old cells of the SSA strain with stable RFP (Fig. S4). The fractional increase in average post-doxycycline budding interval measured for this strain was also reduced compared to the increase in budding intervals measured for the strains containing stable RFP, which is consistent with reduced cell-cycle arrests induced by DSB (Fig. S12). We attribute these observations to the presence of the degron we tagged to the RFP. The degron we used is a truncated Cln2 C-terminal destabilization sequence found in cyclin Cln2. Cln2 is one of three G1 cyclins which together with Cdc28 activate the transcription factor SBF^32^, leading to expression of genes involved in the G1/S transition. Once the cell is out of G1 phase, the expressed S and G2 cyclins activate Cdc28 to promote HDR^26^. The Cln2 C-terminal destabilization sequence encodes protein domains that allow Cln2 to be recognized and targeted for proteasomal degradation by the SCF E3 ubiquitin-ligase complex containing F-box protein Grr1^33^. Given the low abundance of Grr1^33^, constitutive expression of the Cln2 degron tagged to RFP could lead to competition with endogenous Cln2 for recognition by Grr1 and subsequent degradation, which would be expected to result in higher levels of endogenous Cln2. Indeed, deletion of Grr1 has been shown to result in increased Cln2 levels^34^. A higher level of Cln2 in the degron-carrying strain would be expected to speed up the G1/S transition, resulting in a greater percentage of cells with HDR-promoting Clb-Cdc28 activity. This would explain the higher SSA repair efficiency observed in the strain containing the degron-tagged RFP.

Given the above mechanism, and the previously-observed single-cell level correlation between G1 phase duration and total budding interval in old cells^30^, old cells expressing the degron-tagged RFP would be expected to have shorter budding intervals than old cells expressing stable RFP. To assess whether this was the case, we compared average budding intervals in two strains carrying the SSA reporter cassette and lacking the I-SceI cutsite. One strain contained the RFP without the degron tag while the other strain contained the degron-tagged RFP. During the 9-hour post-doxycycline time window, old cells from the two strains displayed similar average budding intervals (Fig. 4d). However, at the time of doxycycline exposure, the average age of the degron-tagged RFP containing strain was two generations greater than that of the stable RFP containing strain (Fig. 4e). To take into consideration this age difference, we next compared average budding intervals between the two strains by using old cells going through more similar ages. For this, we measured the average of all budding intervals occurring after the 15^th^ generation for each cell, and saw that cells of the degron-containing strain had shorter budding intervals (Fig. 4f). At the single-cell level, the right tail of the average budding interval distributions we measured after the 15^th^ generation was longer for the stable RFP containing strain compared to the strain carrying the degron-tagged RFP (Fig. S10). Therefore, the difference in SSA repair efficiency between the two strains can be attributed, at least in part, to the changes in Cln2 levels.

### Assessing age-associated SSA repair efficiency using a reporter with imperfect sequence homology

Homologous recombination is dependent on there being a high degree of homology between the DSB site and repair template. In the context of SSA, strong homology between the direct repeats flanking the DSB is important for repair. Previous work showed that, in the presence of as little as 3% heterology, there can be a six-fold reduction in the ability of cells to complete SSA repair, a phenomenon called heteroduplex rejection^35^. Several mismatch repair proteins and the Sgs1 helicase play an important role in heteroduplex rejection^35^. An age-related change in the rate of heteroduplex rejection would have consequences for the mutation rate in older cells, since heteroduplex rejection prevents recombination between homologous sequences with high levels of sequence heterology.

To determine whether the efficiency of heteroduplex rejection changes with age, we modified the single-strand annealing reporter to have 3% heterology between the non-functional YFP repeats. To assess if the SSA repair sensitivity to this level of heterology changes with age, we measured the rate of YFP repair in young and old cells as before (Fig. S3,f,g, Fig. 5a). As with the strain containing perfect homology (Fig. 2c), the repair efficiency in old cells was lower than it was in young cells (27.5% vs. 34.9%, P-value 0.2787, Fisher’s exact test) (Fig. 5a). Among the young and old cells that were unrepaired at the 9-hour post-doxycycline addition time point, only a small fraction repaired within the next 5 hours (Fig. S5), consistent with robust heteroduplex rejection in cells from both age groups. In fact, despite the expected reduction in the absolute fraction of repaired cells in this strain relative to the strain with perfect homology, the ratio of the repaired-cell fraction in old age to the one in young age (0.789) was very similar to the same ratio measured in the strain with perfect homology (0.766) (Fig. S11). Consistent with the decline in repair efficiency when there is sequence heterology, the average budding time of the 3%-heterology strain in the 9-hour interval after doxycycline addition was increased to 192±4 minutes for old cells and 190±19 minutes for young cells (Fig. 5c). The ratio of post-doxycycline budding intervals between old and young cells of this strain was also reduced relative to the ratio measured in the strain carrying perfect sequence homology in the SSA repair reporter cassette, indicating that the cell-cycle delays experienced by the young cells are not matched by a proportional increase in the delays experienced by the old cells (Fig. S12). These results indicate that there is no replicative age-related decline in heteroduplex rejection in the context of single-strand annealing. The processes normally responsible for heteroduplex rejection work just as well in old cells as they do in young cells.

**Figure 5.**
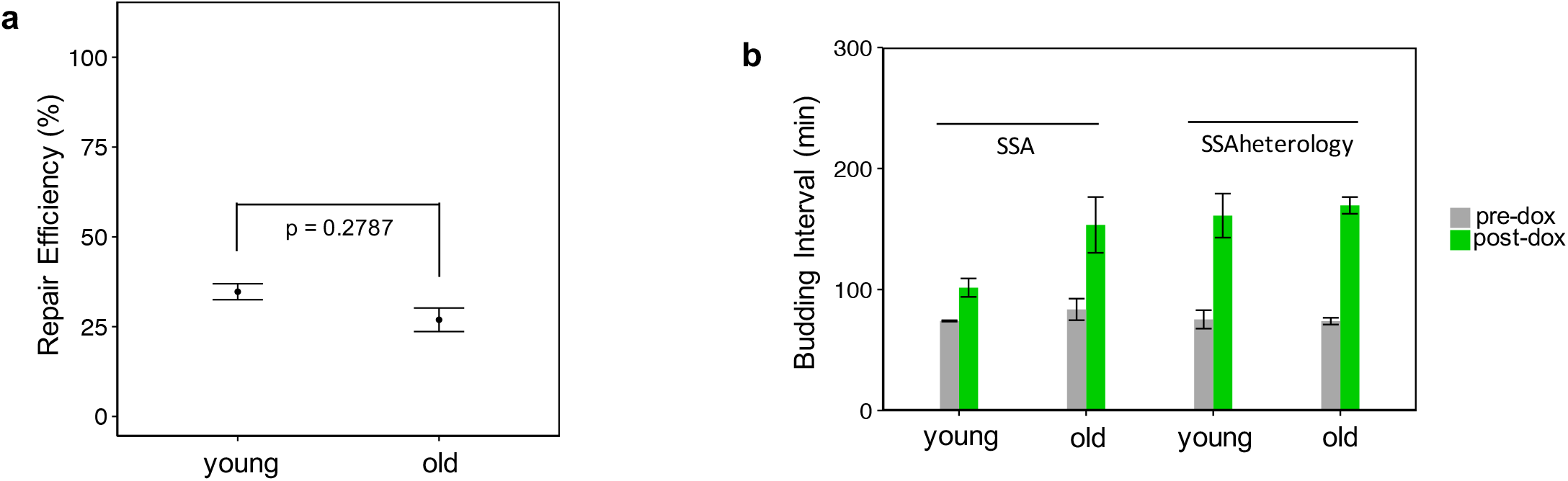
Measuring SSA repair efficiency under 3% sequence heterology. **a.** Old cells show a decline in SSA repair efficiency relative to young cells. Repaired-cell fraction was measured for each replicate to quantify SSA repair efficiency. The mean and SEM (N=2 or 5) of these replicate-based statistics are shown. The replicates (N=5) of the old cells had a mean repair efficiency of 27.5%, while the replicates (N=2) of the young cells had a mean repair efficiency of 34.9%. The p-value of 0.2787 is for a two-sided Fisher’s Exact Test to test for association between repair and age. **b.** Shown are the mean+/-SEM (N=2) of the average budding intervals measured from individual cells of the two replicates during the 4-hour time window before, and the 9-hour time window after doxycycline addition. Cells exhibit an especially large increase in cell-cycle duration after doxycycline treatment (from 75+/-7 min to 161+/-18 min for young cells; from 74+/-3 min to 169+/-7 min for old cells).

## DISCUSSION

Repairing DNA double strand breaks is essential for cells to maintain genome stability and homeostasis. Repair pathway dysfunction can have drastic consequences on a variety of phenotypes in both young and old cells. Despite this, we have very limited understanding about if and how much the efficiency of specific DNA repair pathways change in an age-dependent manner in single cells. The lack of extensive knowledge on this topic is in part due to the historical challenges associated with longitudinally tracking aging cells and simultaneously measuring repair events in them. In this study, using a reporter for detecting single-strand annealing repair in replicatively aging yeast cells, we compared the efficiency of SSA repair for young and old cells, and discovered an age-dependent decline in SSA repair efficiency. Measurement of SSA repair efficiency in strains lacking a key non-homologous end-joining protein or expressing RFP fused to a CLN2-PEST degron provided additional insights into repair pathway usage and efficiency in the context of aging. Finally, we further characterized the change in SSA repair efficiency between young and old cells, in the presence of 3% sequence heterology.

An aging-related decline in the efficiency of single-strand annealing based repair has several important implications. Since the single-strand annealing mechanism shares molecular steps, such as end-resection, with other homology-based repair mechanisms, the molecular reasons behind the reduced SSA efficiency in old cells are expected to affect the efficiency of other homology-based repair mechanisms, as well. The fact that the deletion of the NHEJ protein Dnl4 fails to increase the efficiency of SSA repair suggests that the age-dependent decline in SSA efficiency is not due to increased competition between NHEJ and SSA mechanisms. Given that the repair templates are located close to one another in SSA repair (2.6 kb for the YFP reporter), deficient homolog search cannot easily explain the observed age-related decline in SSA efficiency either. The observation that old cells of a strain containing an SSA repair reporter with RFP fused to a Cln2 degron have higher SSA repair efficiency compared to the degron-free RFP lends support to the mechanism that an increase in G1 duration is partly responsible for the decline in SSA repair efficiency. The single-cell level correlation shown^30^ between the overall cell-cycle duration and G1 phase duration, and our observation of shorter budding intervals for the old cells of the degron-carrying strain, further support the mechanism in which shorter G1 durations are behind the higher repair efficiency measured in the old cells of the degron-carrying strain. Taken together, the age-dependent decline in SSA repair efficiency (Fig. 2c) is due to the old cells’ experiencing longer G1 durations resulting in their inability to activate the steps of the SSA repair mechanism that normally occur outside of G1^10, 15^, such as end-resection, annealing, flap cleavage, gap-filling, or ligation. While our results and the model we propose are novel elucidations of DSB repair in the context of replicative aging of single cells, the broad idea of cell cycle-dependent regulation of DSB repair and the potential role of cyclins had been previously proposed in non-aging contexts^26^.

Our experiments with an SSA strain containing 3% heterology between the direct repeats showed a repair efficiency decline (between young and old cells) similar to the decline observed in the cells of the SSA strain containing perfect homology (Fig. S11). Since heteroduplex rejection in SSA is dependent on both recognition of mismatches by mismatch repair proteins Msh2 and Msh6, and unwinding of the heteroduplex by the Sgs1 helicase, this result implies that these three proteins do not show a significant decline in activity in old cells^35^. Given that an increased mutation rate from recombination between non-allelic sequences would be detrimental to cellular fitness, it is not surprising that heteroduplex rejection does not decline with replicative age. For the cells of the SSA strain containing 3% heterology, the similar drop of repair efficiency in the old cells relative to the young cells is also consistent with a model in which the factor that is responsible for the general decline in SSA efficiency acts independently of heteroduplex rejection.

The relationship between replicative aging and genomic stability has great relevance to our understanding of human health and disease. An increase in genomic instability with age has been hypothesized as a potentially reason for the increased rate of cancer in older individuals^4^. In this study, we used a simple eukaryotic model system to learn whether replicative aging actually coincides with defective DNA repair. We show that replicative aging results in a cell-cycle progression-related decrease in the efficiency of SSA repair, but not heteroduplex rejection. These results raise the possibility that, while old cells show a decline in the efficiency of homology-based repair, they do not show a decline in the efficiency of the mechanisms that prevent aberrant recombination. The age-associated decline we observed in SSA repair efficiency also raises questions about the repair outcome in old cells that are unable to perform SSA repair, since alternative outcomes like MMEJ repair or checkpoint adaptation in the absence of repair^36^, could adversely affect the genome and limit replicative lifespan. Determining what repair outcomes increase in frequency at the expense of SSA and other HDR repair pathways is therefore an exciting research area for future exploration.

## MATERIALS and METHODS

### Construction of plasmids and yeast strains

All strains in this study were constructed in a BY4742 haploid strain background with *MET15* deleted and *LYS2* intact. Table S1 shows the complete strain descriptions and genotypes.

For all SSA repair reporter constructs, the 5’ non-functional mCitrine repeat consists of the 5’ 195 bp of mCitrine followed by a TAG stop codon. The 3’ nonfunctional mCitrine repeat consists of the full 717 bp mCitrine sequence except for the start codon. A 420 bp *TEF1* promoter drives expression of the 5’ non-functional mCitrine, with a 507 bp *TEF1* terminator. A 455 bp segment of DNA from the pRS401 plasmid (lacking homology to the rest of the repair cassette) lies between the *TEF1* terminator and I-SceI cutsite. In total, 1 kb separates the 5’ mCitrine and the I-SceI cutsite. Directly downstream of the I-SceI cut site, a 720 bp *ADH1* promoter drives mCherry with a 207 bp *ADH1* terminator. Immediately downstream of the *ADH1* terminator is the 3’ ATG-less mCitrine followed by a *CYC1* terminator. Altogether, 1.6 kb separates the I-SceI cutsite from the 3’ mCitrine. The *TEF1* and *ADH1* promoters, and *TEF1* and *ADH1* terminators are all from the S288C strain background. While mCitrine and YFP may be used interchangeably throughout the paper, they both refer to mCitrine. Also, while mCherry and RFP may be used interchangeably throughout the paper, they both refer to mCherry.

The mCitrine repeats described in the above paragraph were derived from the yeast codon optimized mCitrine on plasmid pKT0211^37^. pKT0211 was a gift from Kurt Thorn (Addgene plasmid # 8734). The mCherry was also yeast codon optimized and amplified from plasmid pFA6-link-yomCherry-CaURA3^38^. pFA6-link-yomCherry-CaURA3 was a gift from Wendell Lim & Kurt Thorn (Addgene plasmid # 44876).

The SSA control construct is identical to the SSA repair reporter construct described above, except that each 18bp I-SceI sequence has been replaced with the sequence TTGACAATTATTAGTACA. The SSA repair reporter construct carrying 3% heterology is identical to the SSA repair reporter except for 6 wobble position changes in the 5’ non-functional mCitrine. These changes occur at positions 27, 54, 81, 111, 132, and 174.

The mCherry degron (abbreviated ‘RFPdegron’) variants of the SSA reporter constructs contain a 426 bp CLN2-C-terminal destabilization sequence tagged to the C terminus of mCherry. This sequence is a truncation of the CLN2-C-terminal destabilization sequence found in a plasmid containing dsGFP^39^. It encodes amino acids 368-487 of Cln2 immediately followed by amino acids 525-546 of Cln2. The truncated sequence still contains the PEST and D domains of Cln2 that are involved in Cln2 degradation. The plasmid containing dsGFP, PTEF1-yEGFPCLN2PEST-pRS406, was a gift from Claudia Vickers (Addgene plasmid #64406).

All SSA constructs were constructed in pRS306 plasmids carrying the *URA3* marker. The SSA construct was located at the multicloning site with a reverse orientation relative to the *URA3* marker. For integrating the construct, long homologies to the ura3 locus were cloned upstream of the *URA3* and upstream of the SSA reporter (distal to the *URA3* marker). A 1280 bp ura3 upstream homology with a natural BseRI cutsite at the end was cloned upstream of the plasmid’s *URA3* marker. A 669 bp ura3 downstream homology with a natural AatII cutsite at the end was cloned after the P*_TEF1_* driving the 5’ mCitrine repeat. Digestion at the ends of these long homologies using BseRI and AatII generated a linear fragment containing [*URA3* marker + SSA reporter] that was gel-purified and transformed to the *ura3* locus. Integration of the SSA construct was confirmed by colony PCR and sequencing.

Regarding the system of doxycycline-inducible I-SceI expression, a variant of rtTA (rtTA-SEG72P) was used with a less leaky P*_TETO4_* promoter^40^. The yeast strain used to amplify the rtTA variant and P*_TETO4_* were provided by Mads Kaern. I-SceI driven by P*_TETO4_* contains an N terminal NLS and HA tag. The I-SceI DNA sequence was codon optimized for yeast and designed to produce the protein sequence described in previous work^41^.

### Microfluidic chip construction and operation

Detailed information about the PDMS chip construction and its operation is provided in a previous paper from our laboratory^25^.

### Measurement and comparison of age-related changes in SSA repair efficiency

Only cells that were initially YFP^-^ (below threshold) before doxycycline treatment were kept for determination of SSA repair efficiency. Cells obtained from the experiments targeting the old age were required to be >= 15 generations old at the beginning of the doxycycline exposure. Cells obtained from the experiments targeting the young age were required to be <= 5 generations old at the beginning of the doxycycline exposure. Only cells that were alive (un-bursted) and observable at or beyond the 5-hour time window after doxycycline removal were included in analysis. For each experimental replicate, the SSA repair efficiency was calculated as: [(the number of cells repaired within 5 hours after dox removal) / (total number of cells in analysis)].

For each strain, statistical significance of the observed age-related SSA repair efficiency differences was assessed using Fisher’s Exact Test with the R function fisher.test. Single cells from the young cell replicates were pooled together; the same was separately done for the single cells from the old cell replicates. Fisher’s Exact Test was applied to the 2×2 contingency table, with the rows specifying whether the cell was in an old or young SSA repair experiment, and the columns specifying whether the cell was repaired or not repaired based on YFP. The default arguments of the fisher.test function were used, including a two-sided alternative hypothesis.

An estimate for the ratio of old cells’ SSA repair efficiency to young cells’ SSA repair efficiency was obtained by taking the ratio of the old repaired cell fraction to the young repaired cell fraction^42^. Let r_o_ be the number of repaired old cells out of n_o_ old cells, and r_y_ be the number of repaired young cells out of n_y_ young cells. The estimate is then given by the Metric: [(r_o_ / n_o_) / (r_y_ / n_y_)]^42^. Assuming that the log of this metric is approximately normally distributed, we obtain a 95% confidence interval (CI) for it^42^. To calculate the standard error (SE) of the log of the Metric, we assume that the old cell outcomes arise from a binomial distribution with n_o_ trials and the probability of repair pold(repaired)^42^. Similarly, we assume that the young cell outcomes arise from an independent binomial distribution with n_y_ trials and the probability of repair p_young_(repaired)^42^. Then our log(Metric) is an estimator for log(p_old_(repaired) / p_young_(repaired)). Applying the delta method to estimate the variance of our estimator for log(p_old_(repaired) / p_young_(repaired)), we obtain a approximate variance of 1/r_o_ + 1/r_y_ - 1/n_o_ - 1/n_y_^42^ The square root of this approximate variance is the standard error(SE). The 95% CI for the log(Metric) is then given by [(r_o_ / n_o_) / (r_y_ / n_y_)] ±1.96*SE. The error bars on Figure S5 show the antilog of this CI.

### DSB induction experiments in aging cells

To study the effect of replicative age on SSA repair, cells were aged in a microfluidic chip (Fig. 1b) designed to trap virgin mother cells and wash away budded daughter cells. The day before each experiment, a single colony of the strain to be tested was picked to 2 ml CSM 2% glucose and vortexed. 30 *μ*l of the suspended cells was added to 5 ml CSM 2% glucose. After 14 hours of growth in a shaker-incubator (225 rpm) set to 30 C, the culture was diluted and grown further so that at the time of loading the chip, the OD_600_ was between 0.25 and 0.3. For strains carrying an I-SceI cut site, the cultures were also FACS’d to determine which culture had a large fraction of unrepaired, YFP^-^ cells. Immediately prior to loading cells to the microfluidic chip, the cultures were diluted to OD_600_ 0.1. Per lane of the chip, 50-70 *μ*l of the OD_600_ 0.1 culture was loaded at 20 *μ*l/min media flow speed. The base media used throughout the shaker-growth phases and the microfluidic-chip experiments were always CSM minimal media containing 2% glucose as the carbon source. After starting the movie, media flow rate was set to alternate between at 30 *μ*l/min for 2 min and 2.0 *μ*l/min for 8.05 min.

To test SSA repair in young cells, the cells were aged for 7 hours before I-SceI induction. For old cells, the cells were aged for 28 hours before I-SceI induction. Due to differences in cell-division rate, a range of replicative ages are present in the chip before I-SceI induction. Nonetheless, most cells at 7 hours are less than or equal to 5 generations old, and most cells at 28 hours are greater than or equal to 15 generations old (the average age of cells are given in the main text). Only cells in these age ranges are selected for age-based comparison of SSA repair.

After the initial aging period without doxycycline, cells are treated with 10 ug/ml doxycycline media for 4 hours. Doxycyline-free media is restored for the remainder of the movie. SSA repair in each cell is assessed by comparing YFP levels prior to doxycycline treatment and after doxycycline treatment; the assessment is made based on whether or not the YFP level of each cell increases over the threshold level during the 9-hour time period after the addition of doxycycline (4 hours of doxycycline treatment and 5 hours afterwards).

To avoid differences in photo-toxicity prior to I-SceI induction, old and young cells are subjected to the same period of fluorescence snapshots before doxycycline treatment. The fluorescence part of the movie is started 140 minutes before the addition of doxycycline-containing media. YFP/mCitrine (5% Sola intensity, 50 ms exposure time) and mCherry (5% Sola intensity, 200 ms exposure time) snapshots were taken every 30 minutes for the remainder of the movie. PhaseG images were taken every 10 minutes throughout the entire movie. For the degron-tagged mCherry containing strains (referred to as ‘RFPdegron’), the same imaging protocol was used except that for mCherry the Sola intensity was 4% with a 200 ms exposure time.

### Cell-by-cell assessment of YFP repair

After YFP background correction, SSA repair was assessed. A YFP cutoff was first defined, so that cells crossing and remaining above this threshold could be classified as repaired. To determine the cutoff, cells from the SSA control strain were aged to young or old ages (Fig. S1), and imaged by microscopy using the protocol described in the relevant sections above. The mean and standard deviation of YFP levels in the cells of the control strain (not having a cut site) was measured for the 5 time-points prior to dox treatment, to determine the YFP signal due to autofluorescence. This procedure was carried out separately for 2 replicates of young or old cells. The mean + 5*SD was calculated and it less than 6 AU (arbitrary units) for each of the 4 replicates. Based on this, we chose a YFP threshold of 7 AU, values above which indicate the presence of functional YFP.

An R-script was used to identify cells that crossed this 7 AU threshold and remained above it for the remainder of the movie, and the times at which this occurred. The script also recorded cells that crossed and remained above a 300 AU threshold, and the times that this occurred. Cells that crossed and remained above this 300 AU threshold were automatically classified as repaired. Since bright neighboring cells can push the YFP signal of a cell above the 7 AU threshold, cells in which only the 7 AU threshold was crossed were manually inspected movie-by-movie. If no bright neighboring cells were nearby the cell to analyze, the cell was confirmed as repaired. For cells for which the crossing of the threshold was assessed to be likely due to a bright neighbor, the later time points were inspected. If at some later time point, YFP was above the threshold and no bright cells were nearby, then the cell was classified as repaired and this time was recorded. If not, the cell was recorded as unrepaired.

Additionally, the distribution of lag times between crossing of the 7 AU and 300 AU threshold was measured. Based on this distribution, lags > 4 fluorescent measurement durations (120 min) were considered abnormal. For such cells, the crossing of the lower threshold was considered a possible false-positive, and manually inspected to make sure no bright neighbors were nearby. If the crossing of the threshold was due to bright neighbor cells, the later time points were manually scanned to find the first above-threshold YFP measurement that was clearly due to the cell of interest’s internal fluorescence.

### Cell-by-cell assessment of RFP absence in the degron-carrying strain

To determine a cutoff for detecting the absence of mCherry (abbreviated ‘RFP’) expression in cells containing the the degron-tagged RFP within the SSA repair reporter cassette, we measured single-cell RFP levels by quantifying average RFP pixel intensities in YFP^+^ cells. Formation of functional YFP is contingent on loss of the mCherry+degron cassette after cutting. RFP measurements from these cells provide a measure of the intracellular RFP signal as well as cellular autofluorescence and background fluorescence. The RFP cutoff was chosen to be the 95% quantile of the single-cell RFP measurements pooled from the cells that were always observed to be YFP^+^ (Fig. S5). Any RFP expression level below this cutoff was treated as RFP negative while any level above the cutoff was treated as RFP positive.

### Average budding interval measurements

To have a measure of I-SceI DSB-induced cell-cycle arrests, the average budding interval for 3 different time windows was measured: the 4-hour window prior to doxycycline addition (‘pre-dox’), the 4-hour window during doxycycline treatment, and the 5-hour window after doxycycline removal. The average budding interval for the 9-hour time window after doxycycline addition (including the 5-hour time window after doxycycline removal) was also measured. Here, budding interval refers to the difference in time between the appearances of consecutive daughter buds from the same mother cell. For each cell, the budding intervals fully contained within the time window of interest were measured. The full budding interval beginning before, but ending within, the time window of interest was also measured. For each cell and time window of interest, the final partial budding interval was defined as the time between the appearance of the last daughter bud in the time window of interest and the end of that time window. The purpose of measuring this partial budding interval was to be able to detect cell-cycle arrested cells. The average of the final partial budding interval and the full budding intervals measured in the time window of interest was then calculated.

### Image processing

Fluorescence frames were aligned based on non-fluorescent PhaseG images using NIKON NIS Elements software’s image-alignment method, and exported as tifs. A custom Matlab script was used to automatically draw a standard polygon within each trap of the microfluidic chip and save the corresponding binary masks. Masks were manually inspected to make sure the polygons were placed correctly within each trap, so that they would fall within the tracked cells. The masks were applied to the fluorescent images to obtain average mCitrine (‘YFP’) and mCherry (‘RFP’) pixel intensities within each polygon for all time frames in the movie. For situations in which the polygonal region of interest was not fully contained within the cell, a polygon was manually drawn within the cell of interest.

Background fluorescence was subtracted from the cell-specific fluoresence measurements to obtain a more accurate measure of RFP and YFP levels in the cell. Due to spatial and temporal homogeneity in background fluorescence, background fluorescence was obtained for a local, rectangular region around each cell of interest. A custom Matlab GUI was written to identify any non-background region (cell or PDMS) in each rectangle for all time-points of observation. These pixels were excluded in calculating the local background. The average pixel intensity of all background pixels in the local rectangle was defined as the background signal.

A source of error would be present when very bright cells/objects are directly adjacent to the cell of interest. This is not accounted for by the local background pixel average method described above, which includes further away pixels not influenced by the fluorescence of the bright object. To identify these situations, adjacent, non-daughter objects were circled manually using a custom Matlab GUI. If the average pixel intensity of these adjacent objects was greater than 200 AU and if the objects were less than 15 pixels away from the center of the polygonal region of interest, then the local background quantification method described above was not used. Instead, if possible, a test ROI to measure the effect of the bright neighbor object on background was drawn at the same relative distance as the trapped cell ROI. Then, for a given mother cell, the average pixel intensities of these test ROIs was averaged and used as the background to be subtracted.

## ACKNOWLEDGEMENTS

The authors thank Valerie Horsley, Mark Hochstrasser, Weimin Zhong, Ruijie Song, and Acar Laboratory members for useful discussions and feedback on this study. TZY acknowledges support through a Gruber Science Fellowship. TZY was in part supported by a predoctoral training grant from the National Institute of Health (T32 GM007499). MA acknowledges funding from the Ellison Medical Foundation (AG-NS-1015-13) and US National Institutes of Health (1DP2AG050461-01 and 1R01GM127870-01).

## AUTHOR CONTRIBUTIONS

TZY and MA designed the experiments and analysis methods, interpreted the data and results, and prepared the manuscript. TZY constructed the yeast strains, performed the experiments, and analyzed the data. PL prepared the microfluidic chips used throughout the study. MA supervised the study. All authors read and approved the manuscript.

