## Supplementary Information for "Quantitative insights into age-associated DNA-repair inefficiency in single cells"

I. **Supplementary Figures S1-S12**
II. **Supplementary Tables S1-S2**

### I. Supplementary Figures

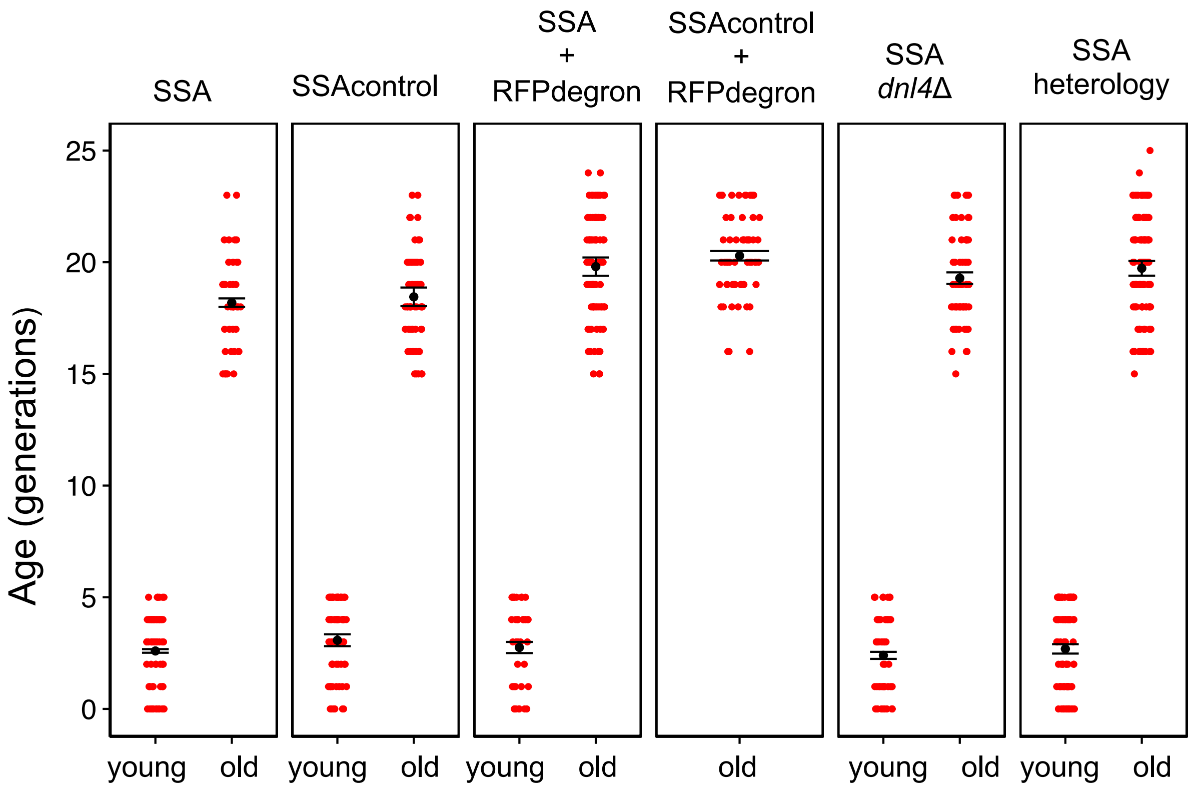


**Figure S1. Ages of cells at the beginning of doxycycline treatment for strains of the study.** Error bars represent Mean+/-SEM of replicative age (after averaging across cells in each replicate) at the beginning of the doxycycline treatment. For both age groups, only cells that were alive 9 hours after doxycycline addition were considered (since these were the only cells used in the repair efficiency calculations). For the old-cell groups, only cells that were at least 15-generations old were included.

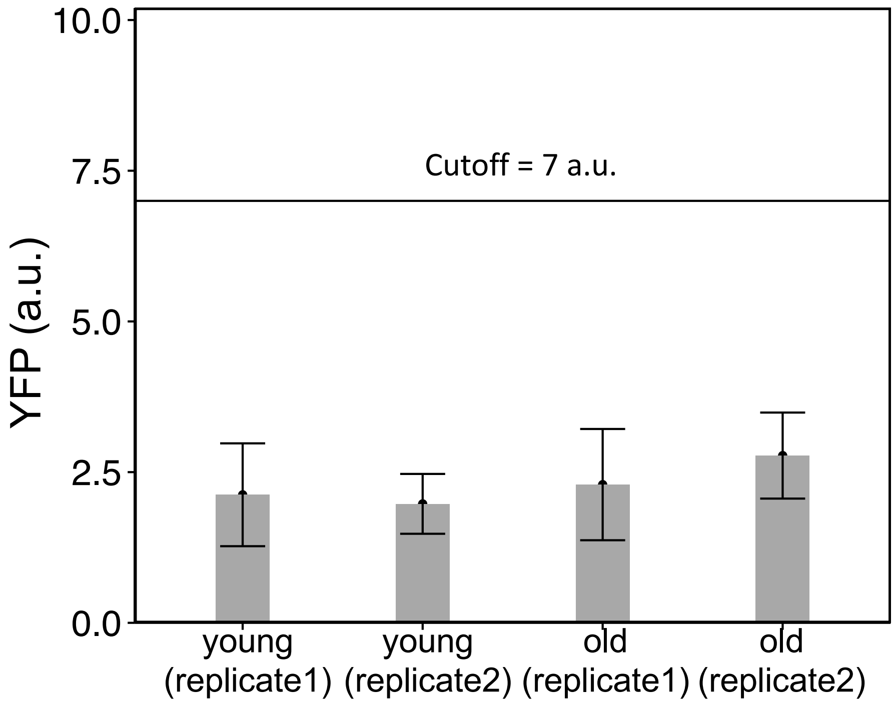


**Figure S2. Determination of the YFP cutoff value for SSA repair based on YFP levels in the SSA control strain without the cutsite.** YFP values were measured for the SSA control strain during the 140 minutes time period (5 fluorescent measurements) prior to doxycycline treatment. Bar heights represent mean of the YFP measurements (in arbitrary units (a.u.)) pooled across all cells of each replicate experiment, and error bars show the standard deviation (SD). The mean + 5SD for each replicate was calculated. Mean + 5SD was 5.4 a.u. and 4.7 a.u. for the young replicates; for the old replicates, it was 6.7 a.u. and 6.4 a.u.. The horizontal line shows the cutoff level (at 7 a.u.) that was used.

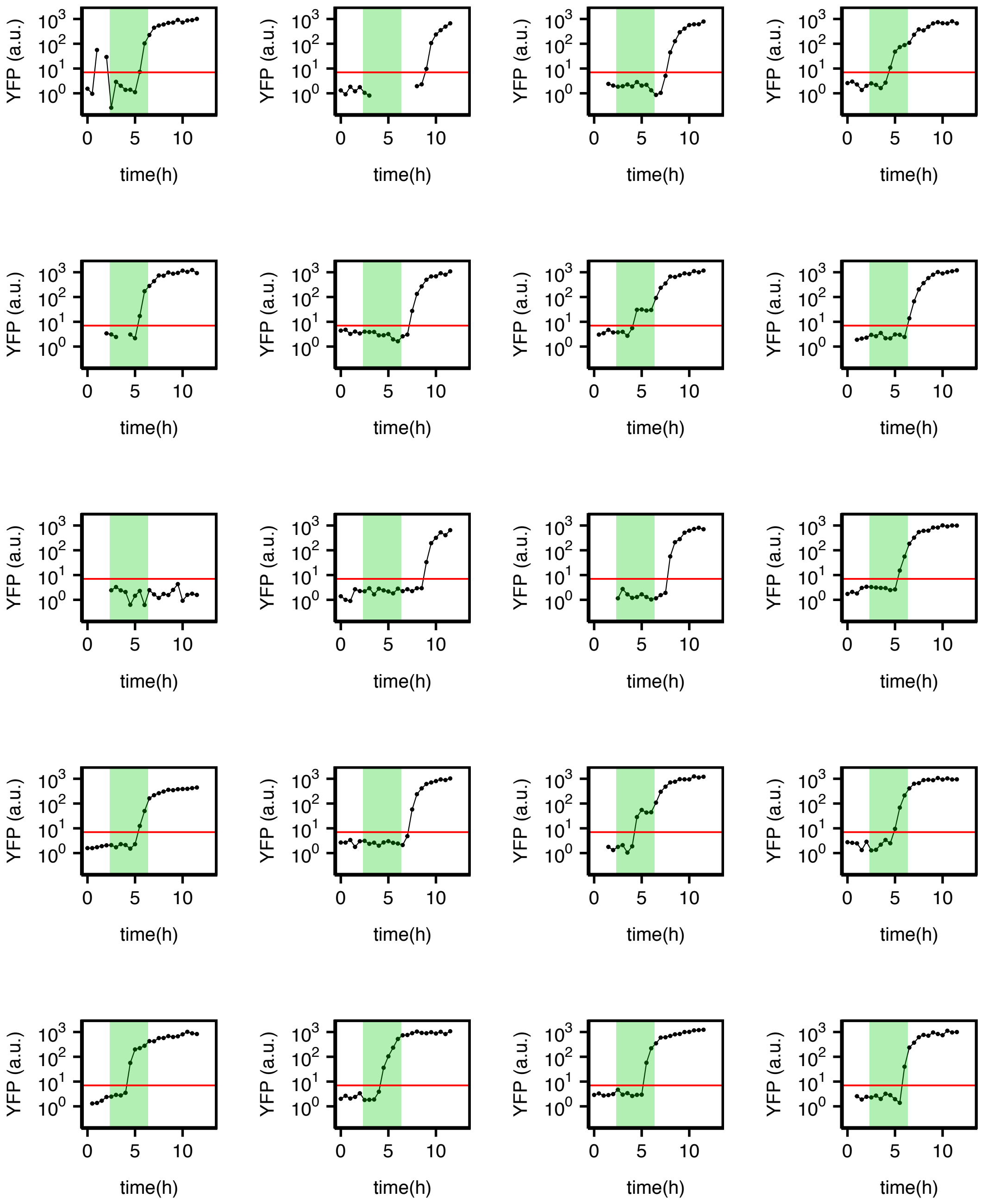


**Figure S3a. Single-cell YFP trajectories for 20 young cells of the strain (yTY125a) containing the SSA repair reporter cassette (replicate #1).** Gaps are due to missing measurements due to inability to measure background fluorescence in the local region around the cell of interest, or negative YFP values which cannot be plotted on the log scale. Shaded green area corresponds to the 4-hour window of doxycycline treatment. Red horizontal line is the YFP cutoff (7 a.u.).

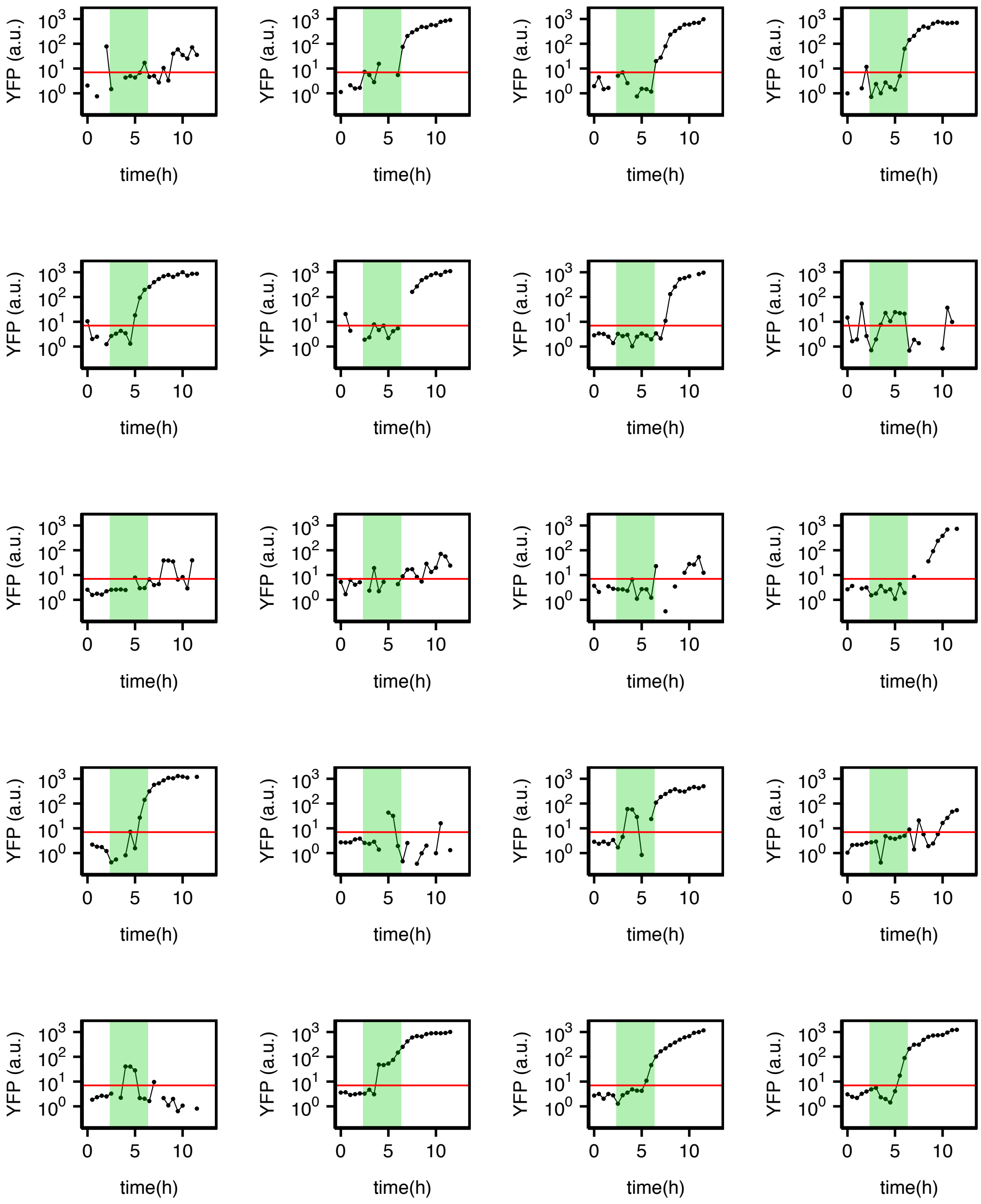


**Figure S3b. Single-cell YFP trajectories for 20 old cells of the strain (yTY125a) containing the SSA repair reporter cassette (Replicate #1).** Gaps are due to missing measurements due to inability to measure background fluorescence in the local region around the cell of interest, or negative YFP values which cannot be plotted on the log scale. Shaded green area corresponds to the 4-hour window of doxycycline treatment. Red horizontal line is the YFP cutoff (7 a.u.).

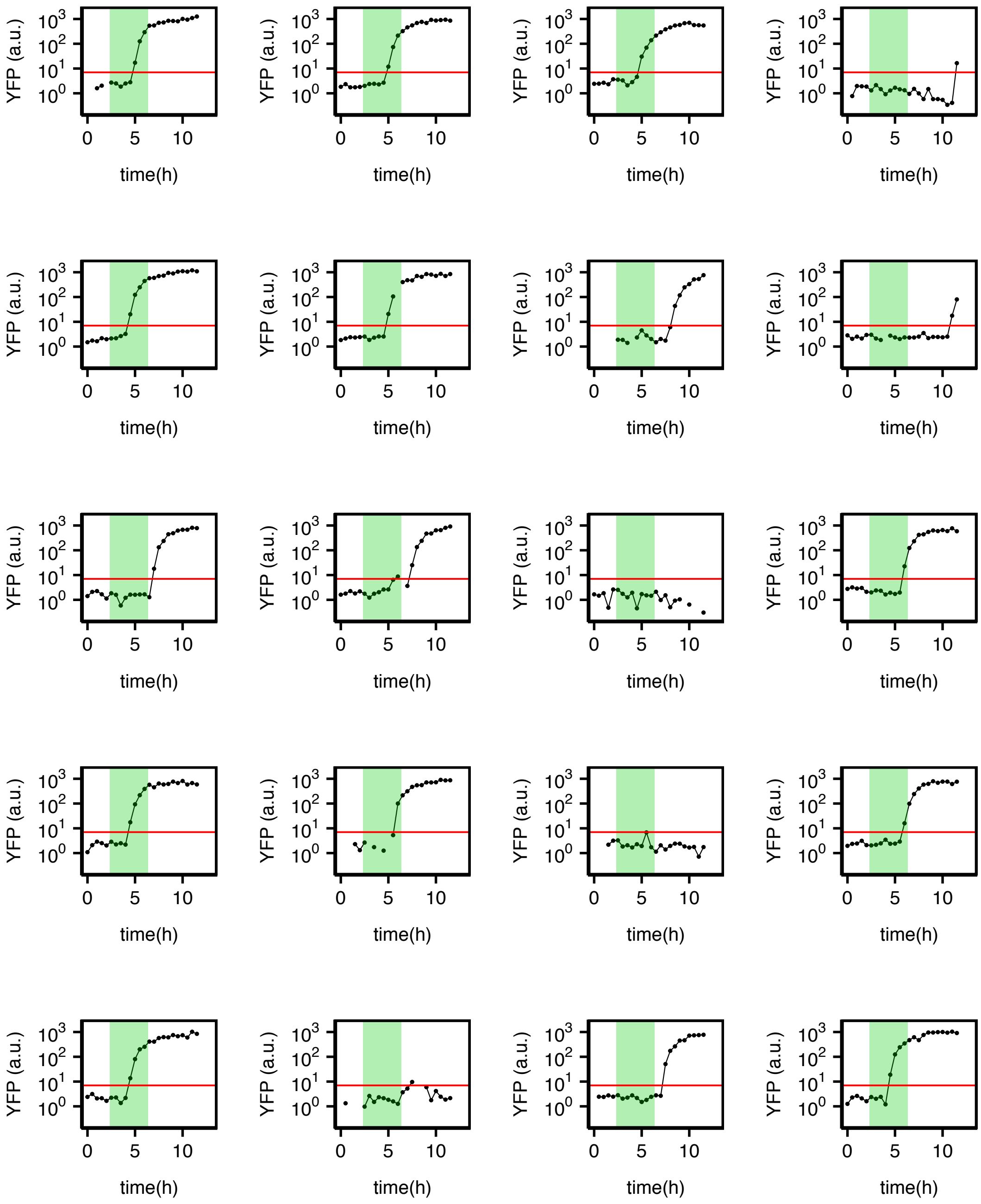


**Figure S3c. Single-cell YFP trajectories for 20 young cells of the SSA strain (yTY147a) containing the SSA repair reporter cassette with the RFPdegron (replicate # 2).** Gaps are due to missing measurements due to inability to measure background fluorescence in the local region around the cell of interest, or negative YFP values which cannot be plotted on the log scale. Shaded green area corresponds to the 4-hour window of doxycycline treatment. Red horizontal line is the YFP cutoff (7 a.u.).

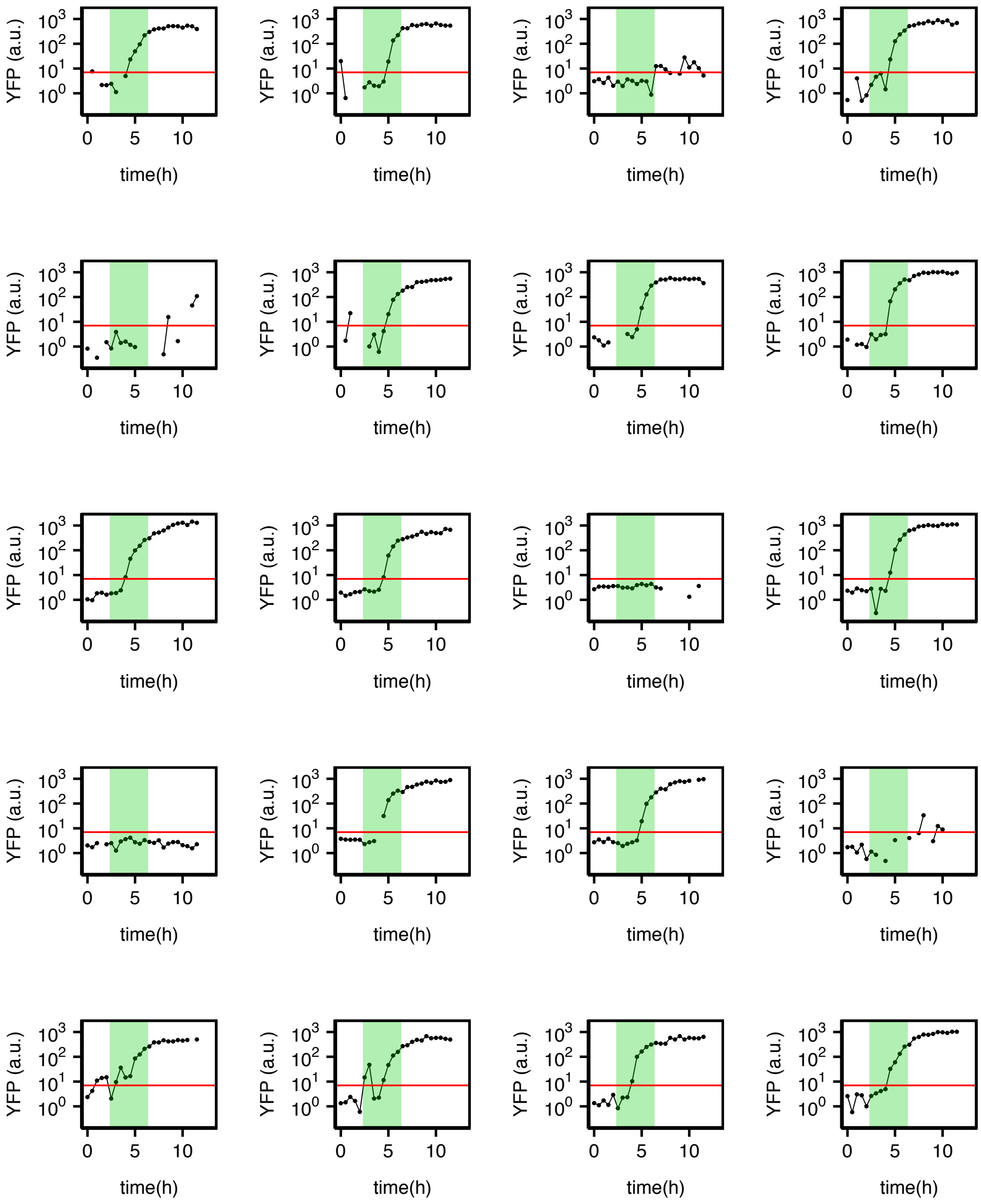


**Figure S3d. Single-cell YFP trajectories for 20 old cells of the SSA strain (yTY147a) containing the SSA repair reporter cassette with the RFPdegron (replicate #1).** Gaps are due to missing measurements due to inability to measure background fluorescence in the local region around the cell of interest, or negative YFP values which cannot be plotted on the log scale. Shaded green area corresponds to the 4-hour window of doxycycline treatment. Red horizontal line is the YFP cutoff (7 a.u.).

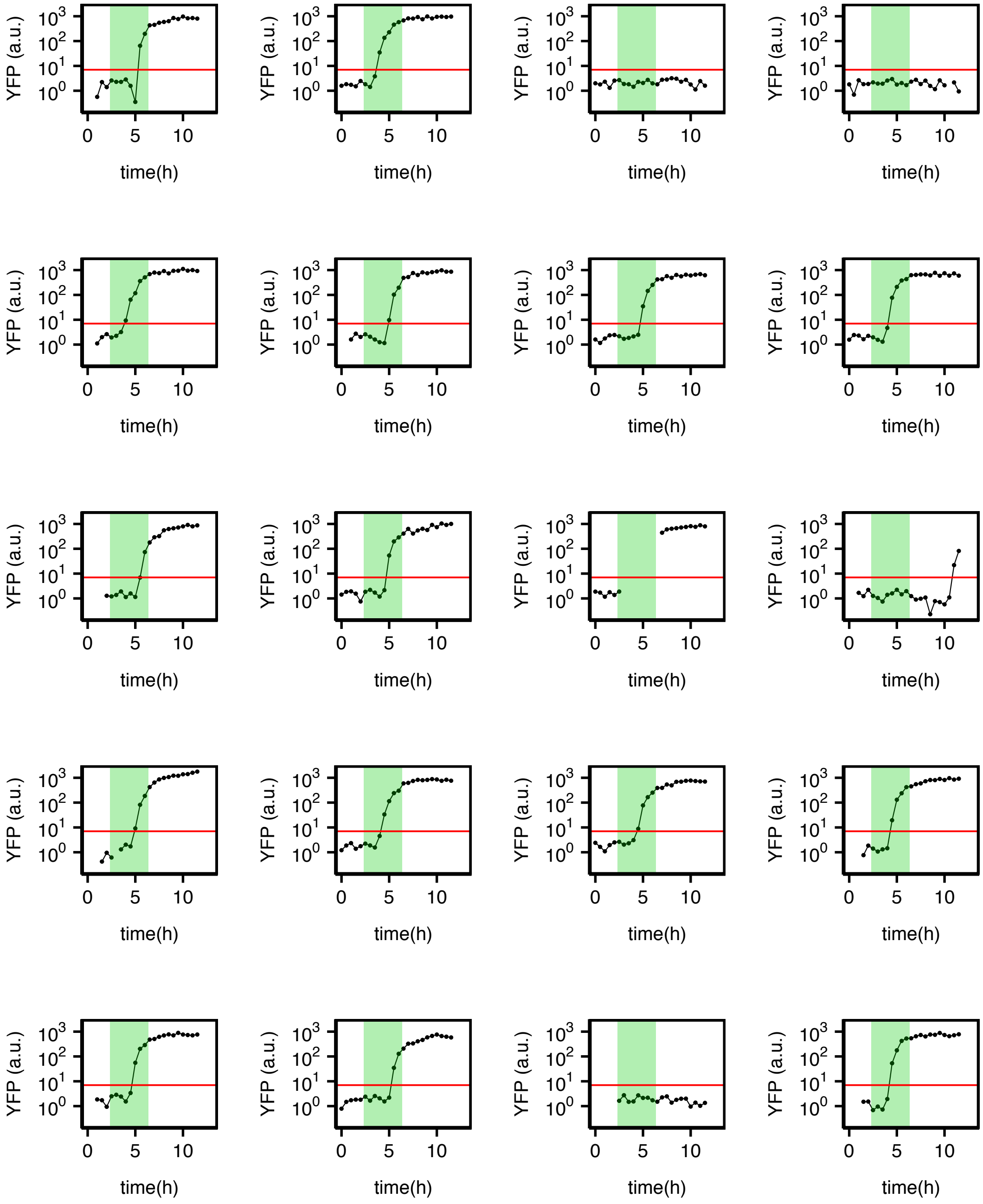


**Figure S3e. Single-cell YFP trajectories for 20 young cells of the *DNL4*-deleted SSA strain (yTY149a) containing the SSA repair reporter cassette (replicate #2).** Gaps are due to missing measurements due to inability to measure background fluorescence in the local region around the cell of interest, or negative YFP values which cannot be plotted on the log scale. Shaded green area corresponds to the 4-hour window of doxycycline treatment. Red horizontal line is the YFP cutoff (7 a.u.).

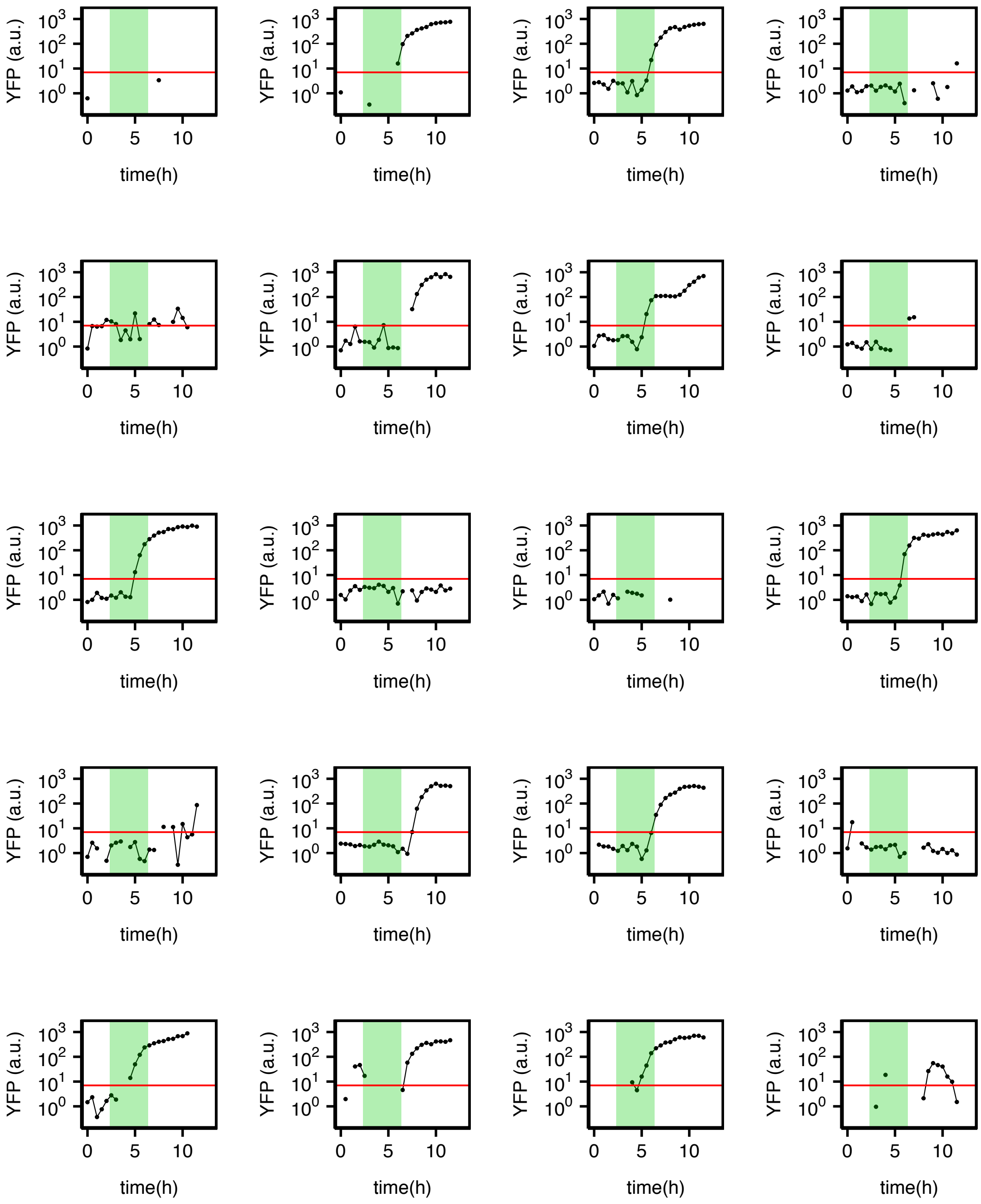

**Figure S3f. Single-cell YFP trajectories for 20 old cells of the *DNL4*-deleted SSA strain (yTY149a) containing the SSA repair reporter cassette (replicate #1).** Gaps are due to missing measurements due to inability to measure background fluorescence in the local region around the cell of interest, or negative YFP values which cannot be plotted on the log scale. Shaded green area corresponds to the 4-hour window of doxycycline treatment. Red horizontal line is the YFP cutoff (7 a.u.).

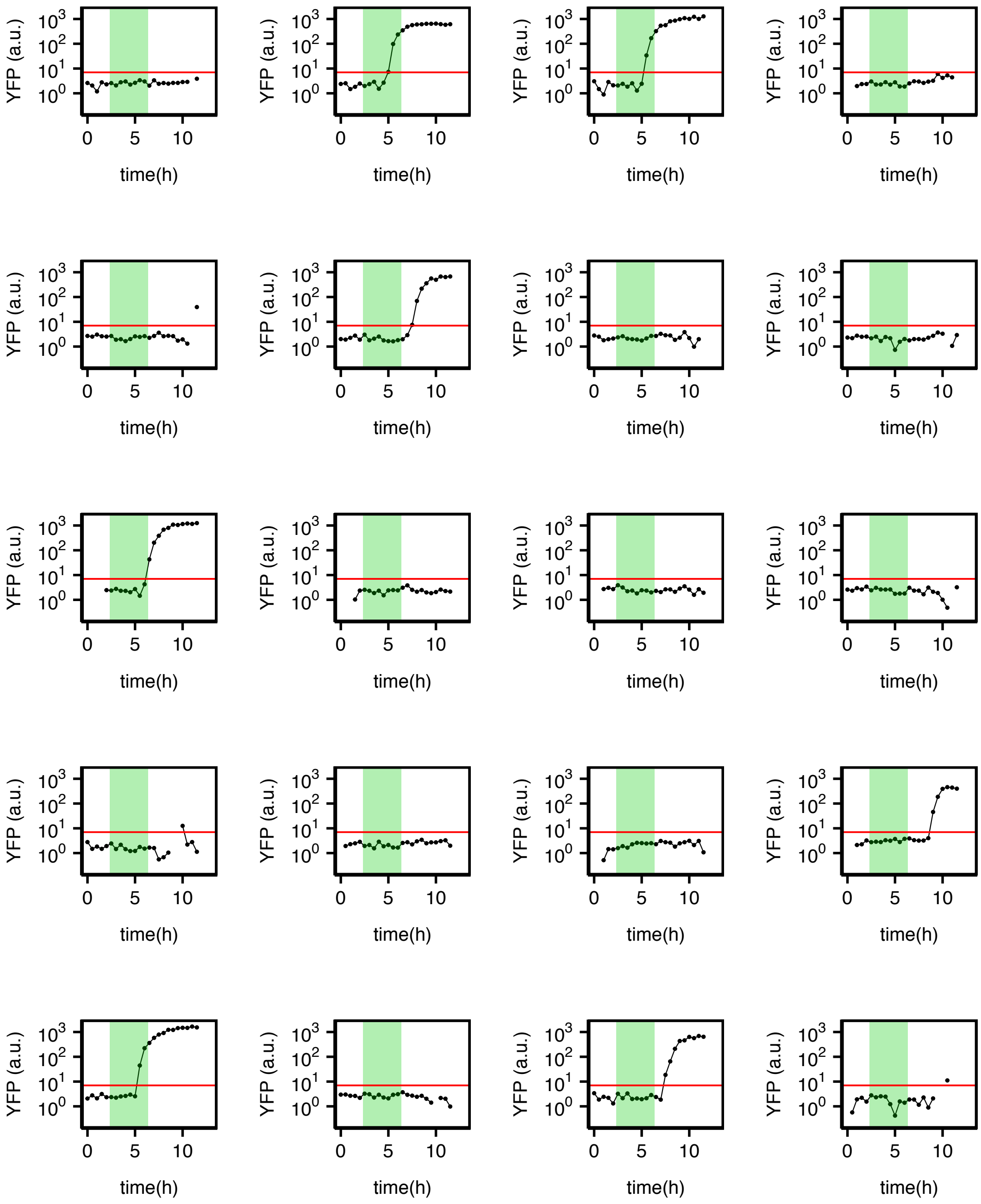


**Figure S3g. Single-cell YFP trajectories for 20 young cells of SSAheterology strain (yTY133c) containing the SSA repair reporter cassette with 3% heterology (replicate #1).** Gaps are due to missing measurements due to inability to measure background fluorescence in the local region around the cell of interest, or negative YFP values which cannot be plotted on the log scale. Shaded green area corresponds to the 4-hour window of doxycycline treatment. Red horizontal line is the YFP cutoff (7 a.u.).

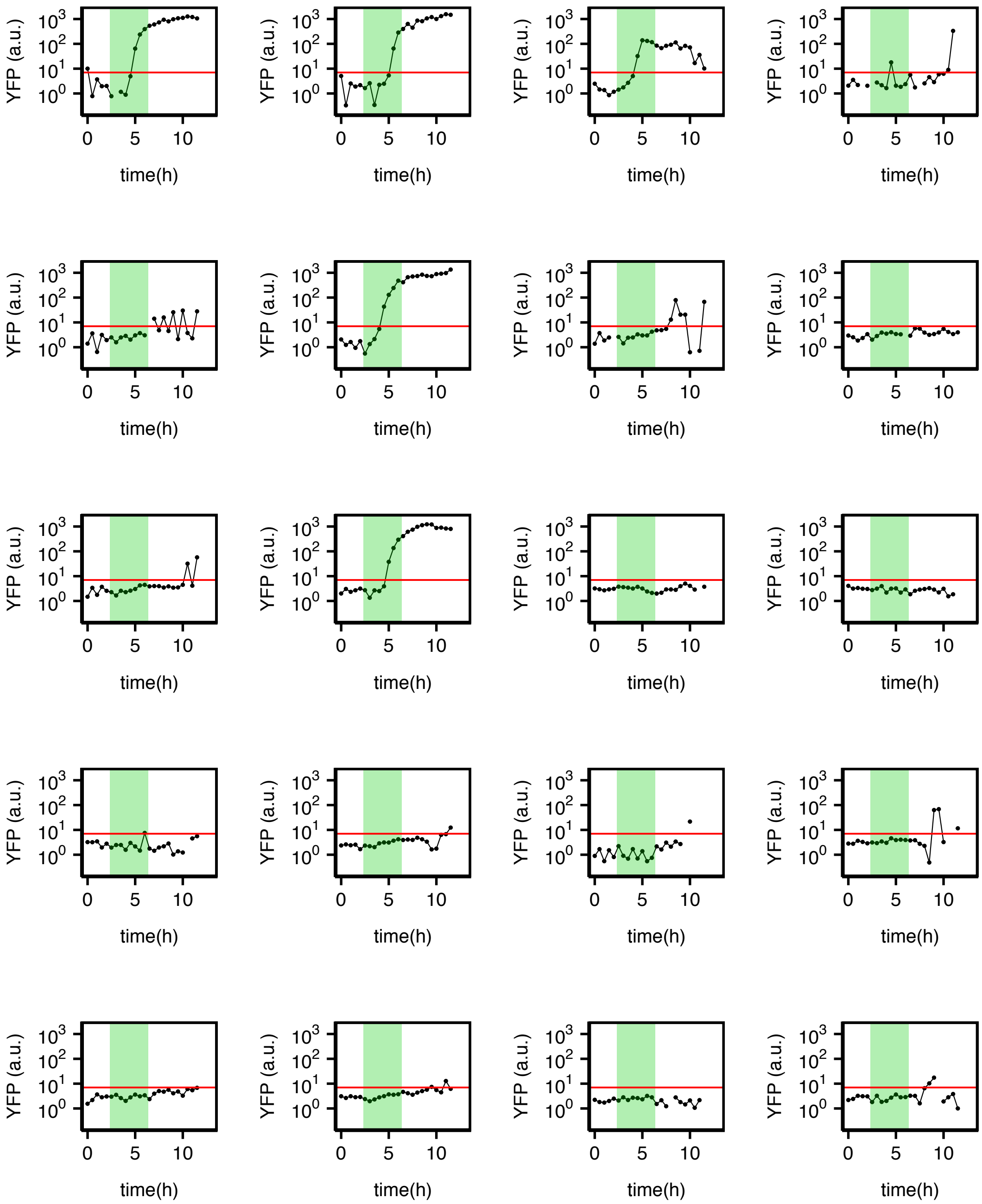


**Figure S3g. Single-cell YFP trajectories for 20 old cells of the SSAheterology strain (yTY133c) containing the SSA repair reporter cassette with 3% heterology (replicate #2).** Gaps are due to missing measurements due to inability to measure background fluorescence in the local region around the cell of interest, or negative YFP values which cannot be plotted on the log scale. Shaded green area corresponds to the 4-hour window of doxycycline treatment. Red horizontal line is the YFP cutoff (7 a.u.).

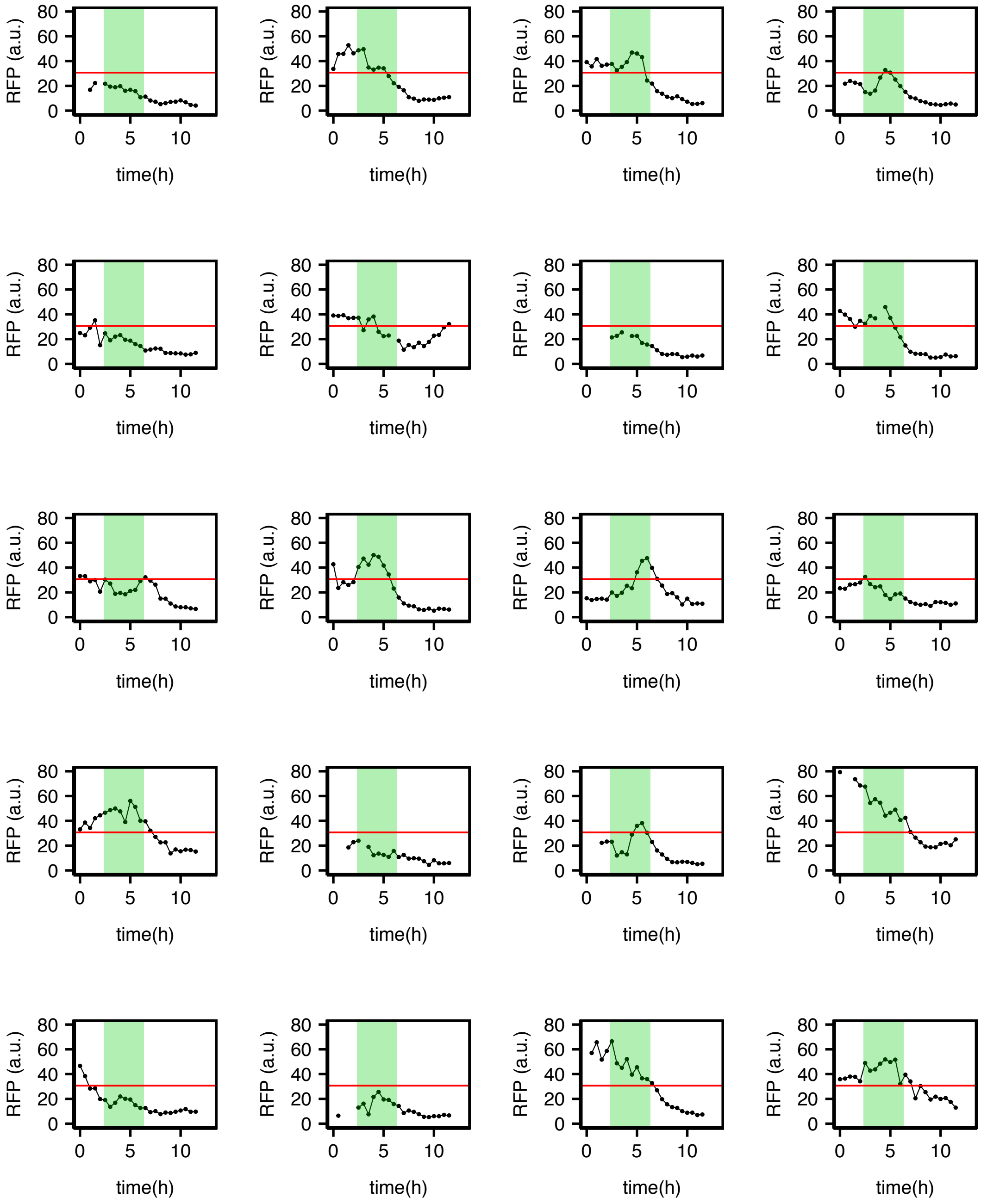


**Figure S3h. Single-cell RFP trajectories for 20 young cells of the SSA strain (yTY147a) containing the SSA repair reporter cassette with the RFPdegron (replicate # 2).** Gaps are due to missing measurements due to inability to measure background fluorescence in the local region around the cell of interest. Shaded green area corresponds to the 4-hour window of doxycycline treatment. Red horizontal line is the RFP cutoff for calling RFP absence.

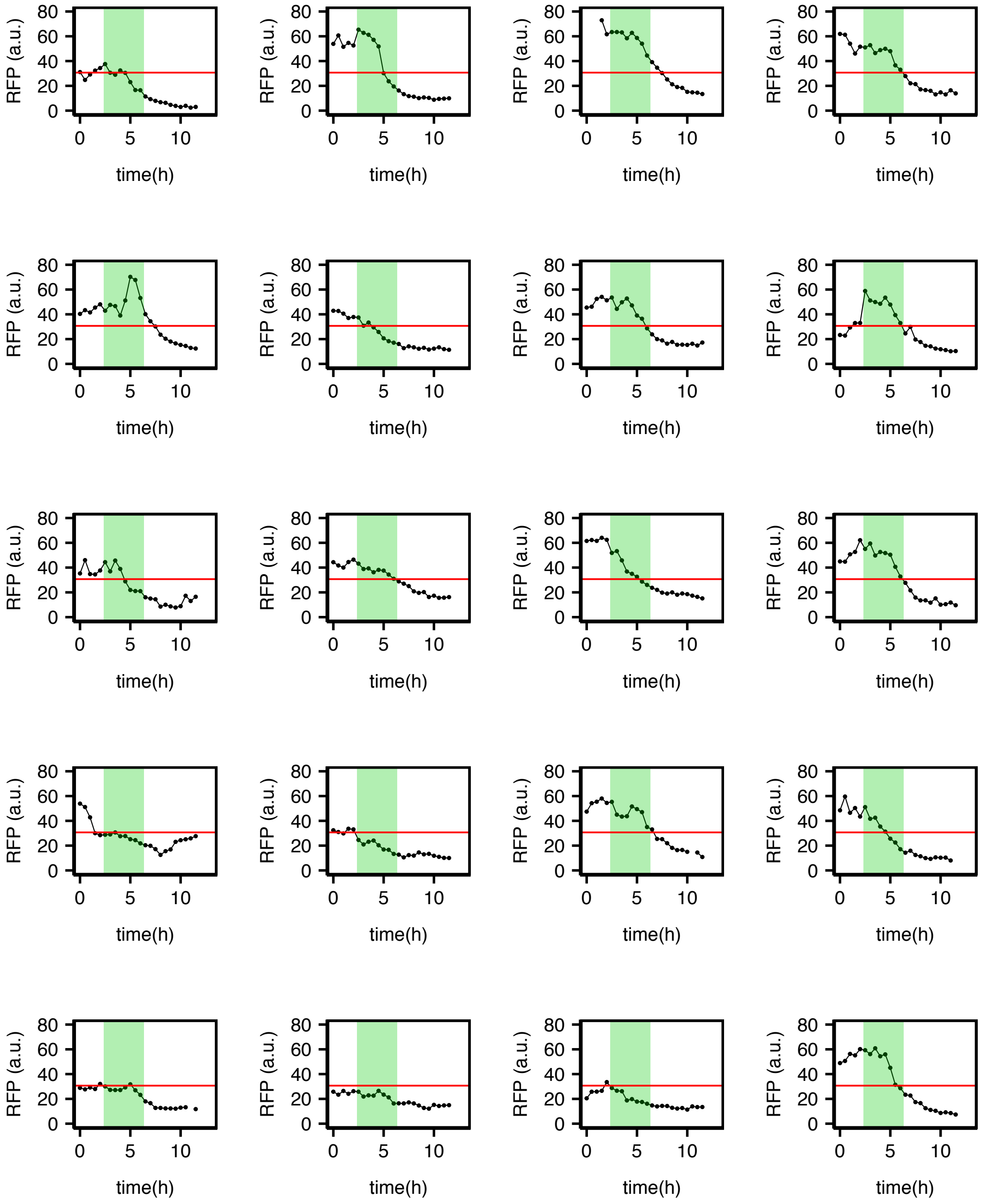


**Figure S3i. Single-cell RFP trajectories for 20 old cells of the SSA strain (yTY147a) containing the SSA repair reporter cassette with the RFPdegron (replicate #1).** Gaps are due to missing measurements due to inability to measure background fluorescence in the local region around the cell of interest. Shaded green area corresponds to 4-hour window of doxycycline treatment. Red horizontal line is the RFP cutoff for calling RFP absence.

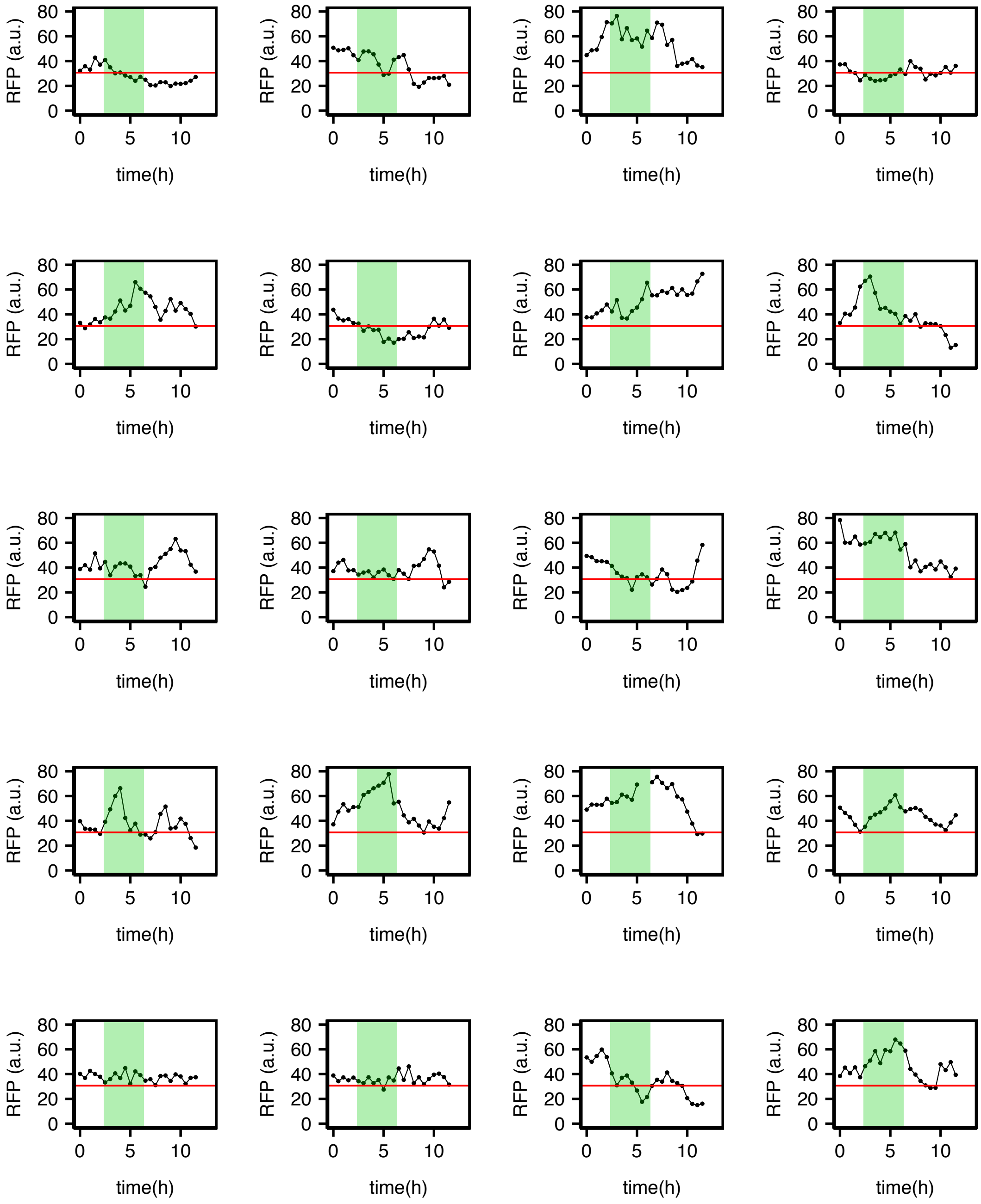


**Figure S3j. Single-cell RFP trajectories for 20 old cells of the SSAcontrol strain (yTY146a) containing the SSA repair reporter cassette with the RFPdegron (replicate #1).** Gaps are due to missing measurements due to inability to measure background fluorescence in the local region around the cell of interest. Shaded green area corresponds to the 4-hour window of doxycycline treatment. Red horizontal line is the RFP cutoff for calling RFP absence.

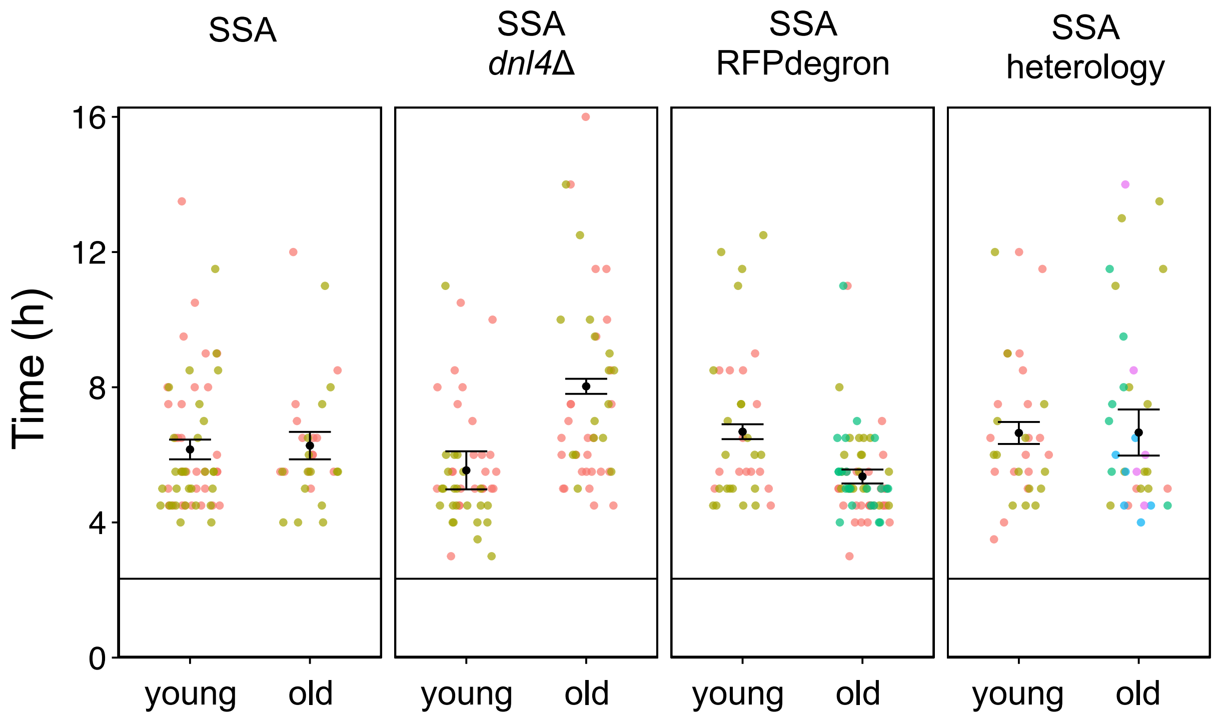


**Figure S4. YFP appearance times in the strains carrying the SSA repair reporter and its variants.** The y-axis indicates the time passed since the start of the fluorescence measurements in each movie. The black horizontal line with y-intercept at 2.3 h corresponds to the time of doxycycline addition relative to the start of the fluorescence measurements. Points correspond to the time of first YFP appearance for individual cells. Error bars correspond to Mean+/-SEM (N=3 for old SSA+RFPdegron, N=5 for old SSA heterology, N=2 for all other strain/agegroup pairs) of the replicate averages (after averaging all cells' YFP appearance times in a replicate). Different colors correspond to different replicates.

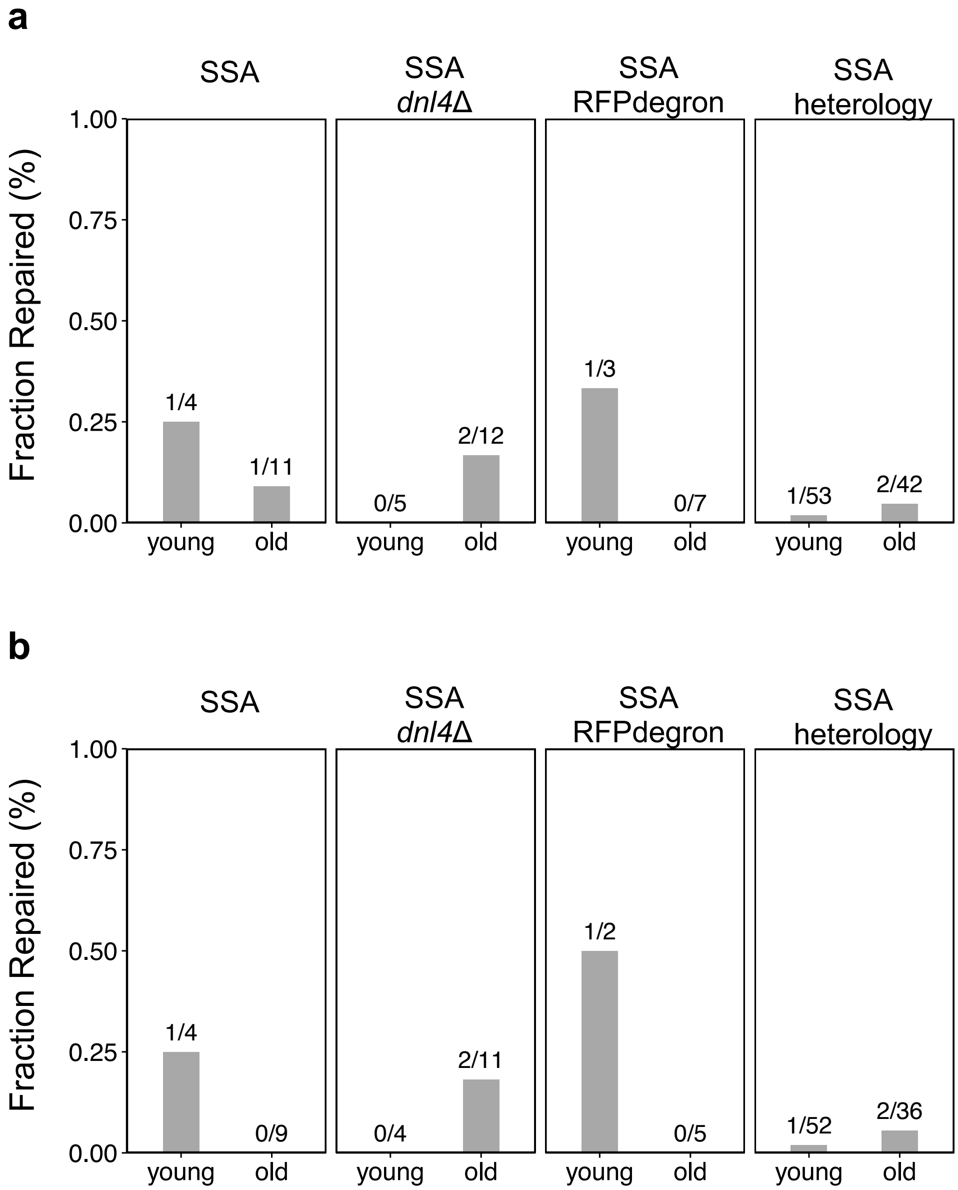


**Figure S5. Assessing SSA repair during the next 5 hours after the 9-hours post-doxycycline addition time window (only for cells unrepaired earlier).** y-axis is the fraction of cells that were SSA unrepaired ($YFP^-$) at the $9^{th}$-hour post-doxycycline addition time point but were are repaired within the next 5 hours (between $9^{th}$ hour and $14^{th}$ hour time points after doxycycline addition). The cell counts used to compute these fractions are shown above the bars. **a.** Fraction of repair computed among cells that either burst in the 9h-14h time window, or were alive at the end of this window. **b.** Fraction of repair computed only among the cells that were alive at the end of the 9h-14h time window.

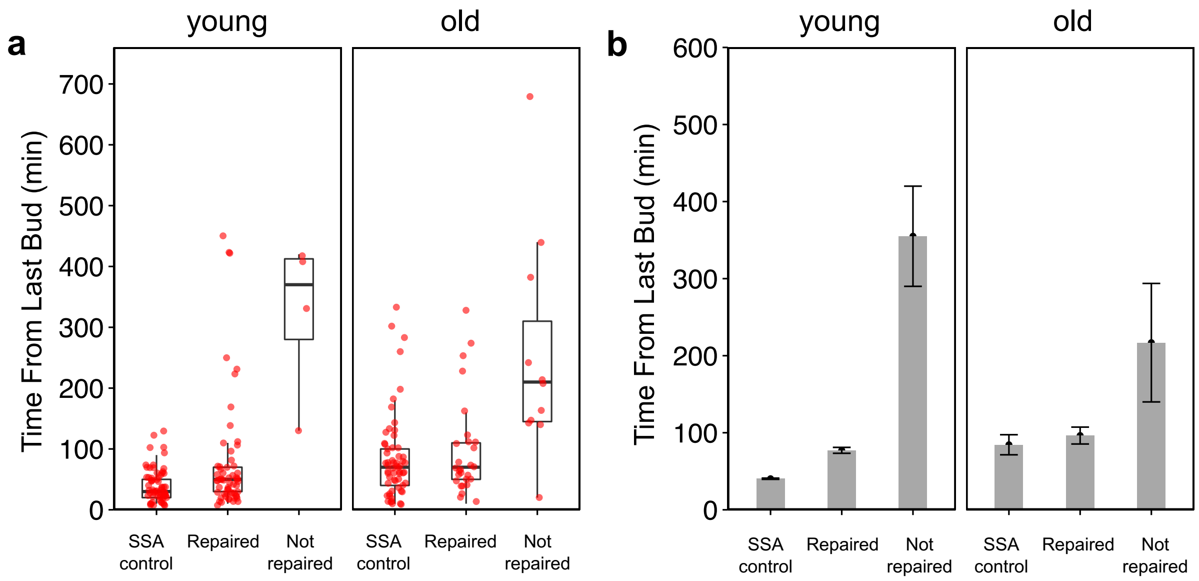


**Figure S6. Time from last bud measured at the 9th hour after doxycycline addition time point for cells carrying the SSA repair reporter cassette.** For each age group (young vs old), cells are grouped by whether YFP was repaired at the end of the 9-hour post-doxycycline time window. Values for the SSA control strain are also shown for comparison. **a.** Values for individual cells from two replicates for each age group. Boxplots are overlaid to show the distribution of times for cells pooled across 2 replicates. **b.** Mean+/-SEM (N=2) of average times from last bud (after averaging across all cells' times from last bud within each replicate). For both age groups, the unrepaired cells have a greater time from last bud at the 9th hour after doxycycline addition compared to the repaired cells (217+/-77 min for unrepaired cells vs 96+/-11 min for the repaired cells).

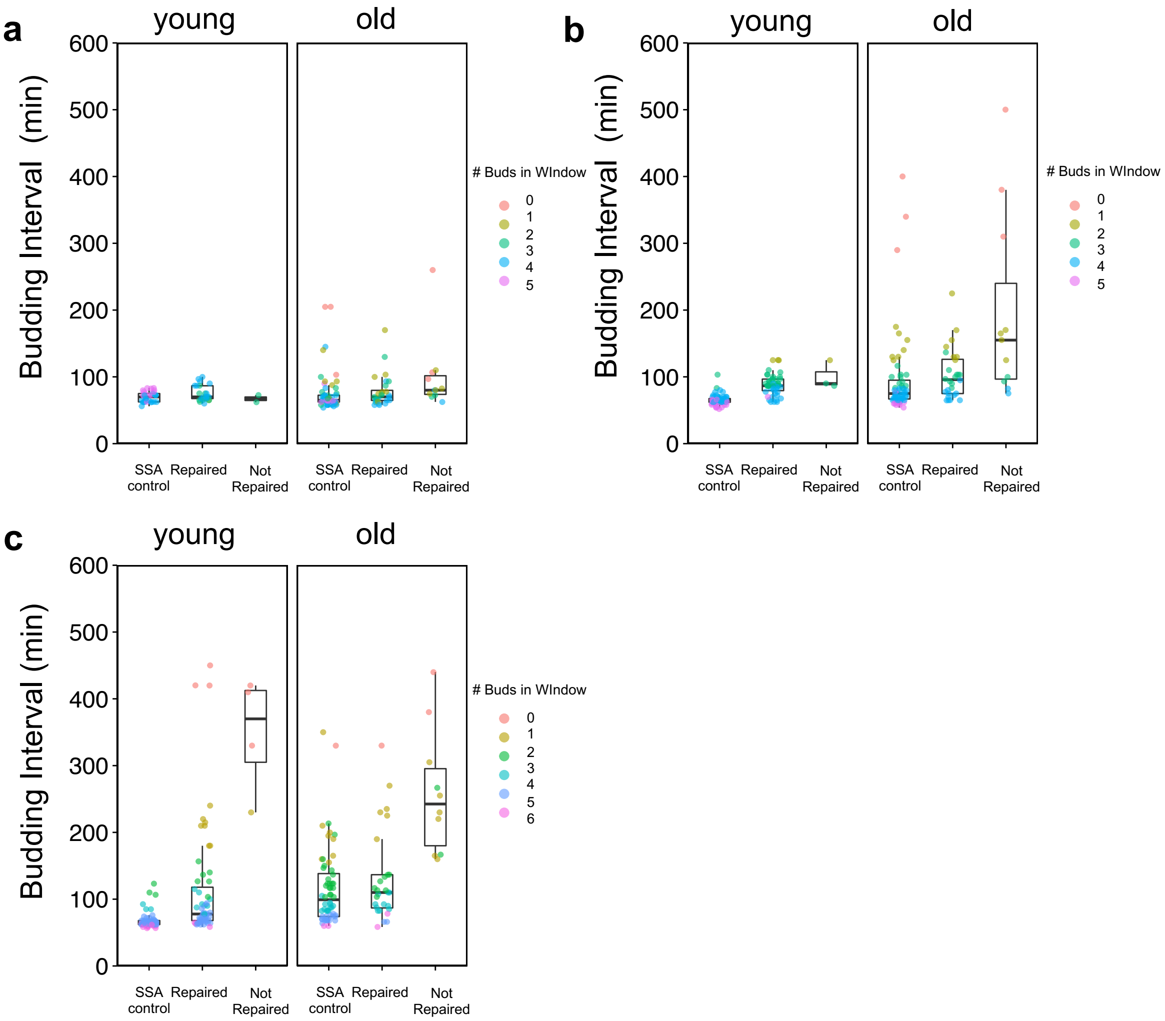


**Figure S7. Average budding intervals before (a), during (b), and after (c) the doxycycline treatment for repaired and unrepaired cells of the strain (yTY125a) containing the SSA repair reporter cassette.** Individual data points correspond to average budding intervals for single cells. Single cell data pooled from 2 replicate experiments is shown, with boxplots for the distribution of values in each sub-group overlaid. **a.** During the 4-hour time window before doxycycline addition. **b.** During the 4-hour time window coinciding with doxycycline treatment. **c.** During the 5-hour time window after doxycycline removal.

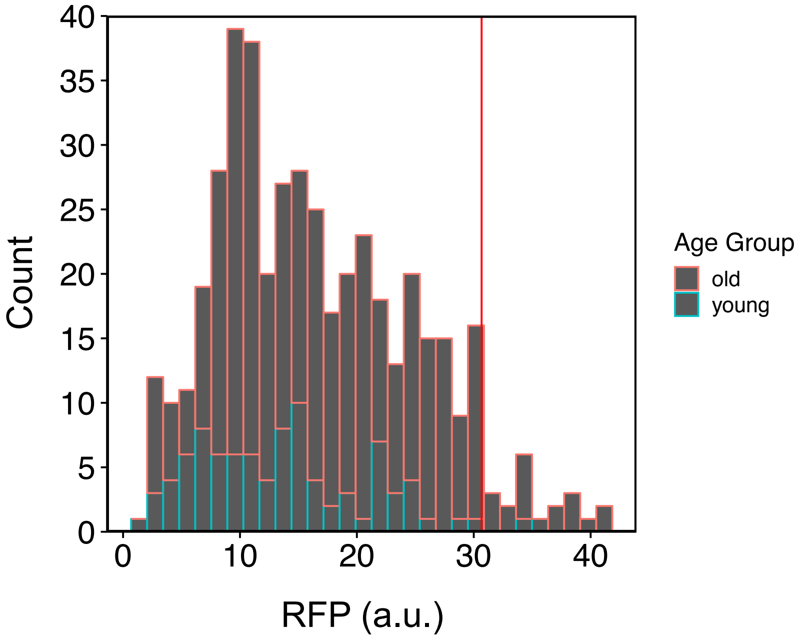


**Figure S8. Defining an RFP cutoff using RFP values measured from the YFP+ cells of the SSA+RFPdegron strain (yTY147a).** The histogram shows the distribution of all RFP measurements over the first 12 hours of the movie, for cells that initially were strongly YFP+ (YFP > 300 a.u.). Bars are color coded to show the counts of measurements from each age group. The red vertical line corresponds to the 95% quantile of this distribution.

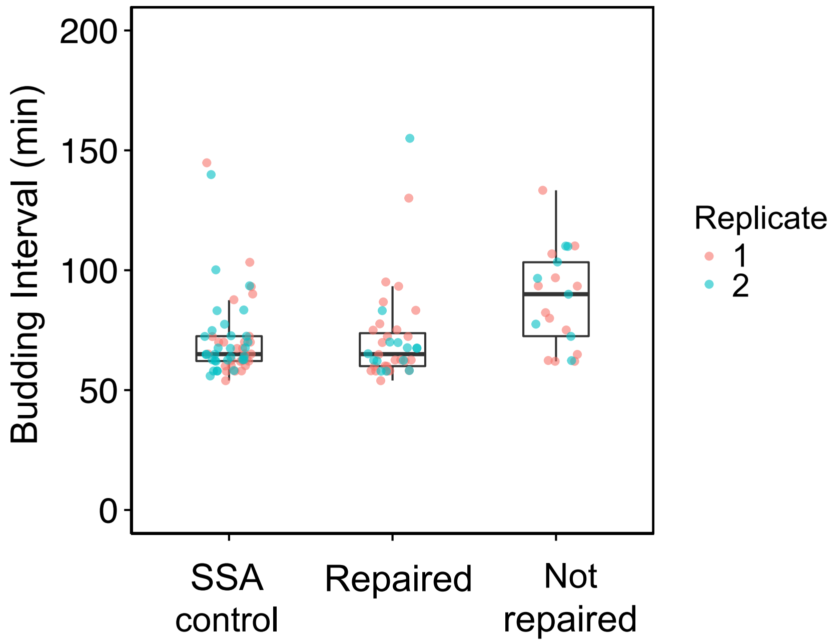


**Figure S9. Average budding interval in the 4-hour pre-doxycycline time window for old cells of the strain containing the SSA reporter but lacking *DNL4*.** Individual points correspond to average budding intervals during the 4-hour pre-doxycycline time window for single cells. Cells are grouped by whether SSA repair was detected by YFP within the 9-hour time window after doxycycline addition. Budding intervals for the control strain lacking a cut-site are also shown for comparison. Boxplots summarizing the distribution of the corresponding single-cell values are also shown.

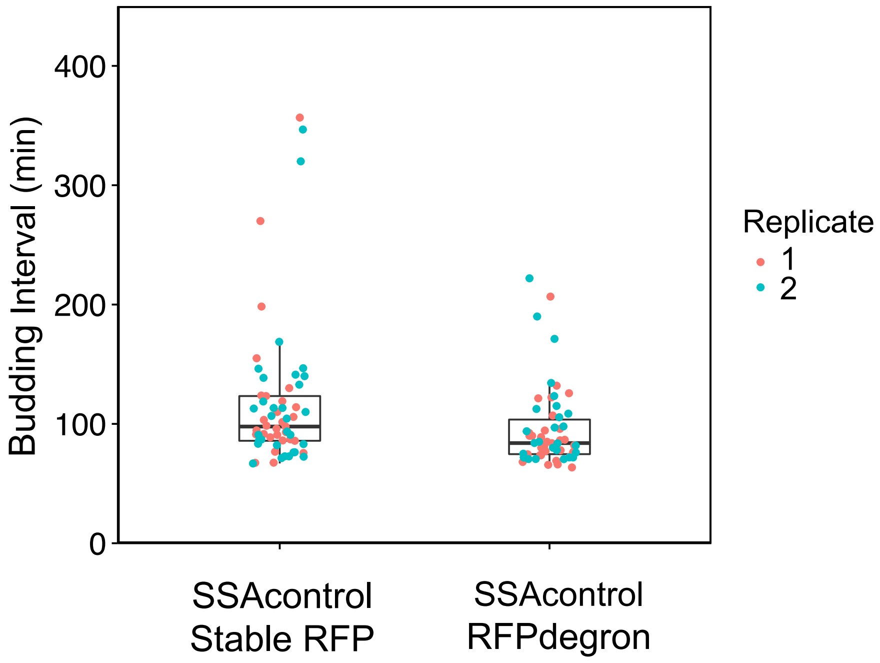


**Figure S10. Average budding intervals measured after doxycycline addition for old cells of the SSAcontrol strains missing the cutsite, with stable RFP (yTY126a) or degron-tagged RFP (yTY146a).** Only cells of age 15 or older at the time of dox addition were considered. Points corresponding to average budding intervals (averaged for each single cell) during this time window are color coded by replicate. Boxplots for the distribution of the single cell values (pooled across the 2 replicates) are overlaid.

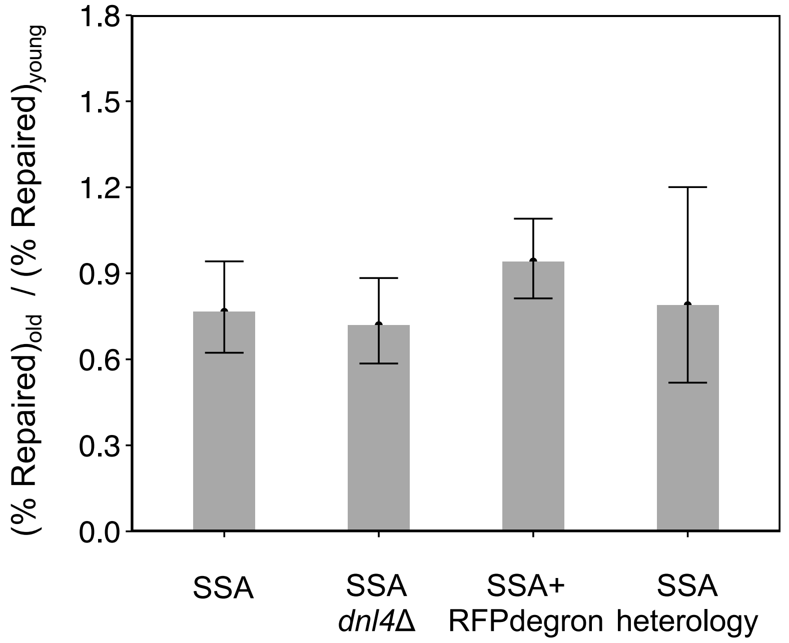


**Figure S11. Ratio of SSA repair efficiency in old cells to young cells of each strain.** Repair efficiency was defined as the fraction of cells repaired by SSA by the end of the 9-hours time window after doxycycline addition, so it is partly a measure of SSA repair speed. For each strain shown, the ratio of the old cell repair efficiency to the young cell repair efficiency was taken. The error bars represent 95% confidence intervals based on applying the delta method to the log ratio of the pooled old cells' repaired fraction to the pooled young cells' repaired fraction (Materials and Methods). The mean SSA repair efficiency ratio is greater for the SSA+RFPdegron strain compared to the other three strains. The specific mean SSA repair efficiency ratios are: 0.941 for the SSA+RFP degron strain (yTY147a), 0.766 for the SSA strain without the degron (yTY125a), 0.719 for the SSA strain lacking *DNL4* (yTY149a), and 0.789 for the SSA strain with 3% heterology (yTY133c).

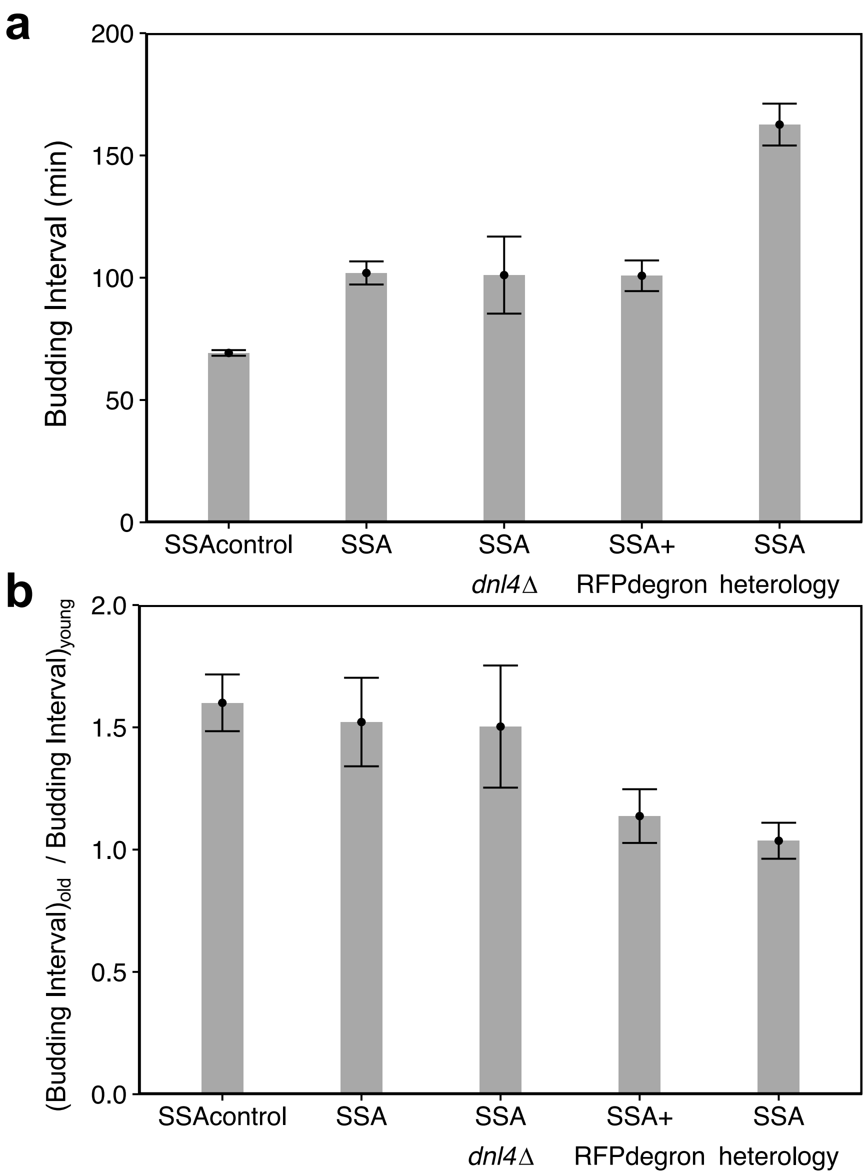


**Figure S12. Strain-wise comparison of age-related slowdown in budding intervals during the 9-hour time window after doxycycline addition. a.** Average budding intervals during the 9-hour time window after doxycycline addition for young cells of each strain. Mean+/-SEM (N=2) of replicate-averaged single cell values for the time window are shown. **b.** Ratio of budding intervals between old and young cells of each strain, measured after doxycycline addition. For each strain, the mean of replicate-averaged single-cell budding intervals was calculated for old-cell replicates and for young-cell replicates. Bar height corresponds to the ratio of these means. Error bars are calculated by propagating the uncertainty (SEM, N=2) of each of the means.

#### II. <u>Supplementary Tables</u>

| Strain | Description | Genotype |
|---|---|---|
| yTY126a ('SSAcontrol' in figures) | SSA (control) | *MATα, his3Δ, LYS2, met15Δ, leu2Δ::LEU2-*$P_{MYO2}$*-rtTA-Adh1t, ho::HIS5-*$P_{TETO4}$*-I-SceI-Cyc1t, ura3Δ::URA3-*SSAreporter(control, no cut-site) |
| yTY125a ('SSA' in figures) | SSA (perfect homology) | *MATα, his3Δ, LYS2, met15Δ, leu2Δ::LEU2-*$P_{MYO2}$*-rtTA-Adh1t, ho::HIS5-*$P_{TETO4}$*-I-SceI-Cyc1t, ura3Δ::URA3-*SSAreporter (perfect homology) |
| yTY133c ('SSAheterology' in figures) | SSA (3% heterology) | *MATα, his3Δ, LYS2, met15Δ, leu2Δ::LEU2-*$P_{MYO2}$*-rtTA-Adh1t, ho::HIS5-*$P_{TETO4}$*-I-SceI-Cyc1t, ura3Δ::URA3-*SSAreporter(with 3% heterology) |
| yTY146a ('SSAcontrol+ RFPdegron' in figures) | SSA (control, RFPdegron) | *MATα, his3Δ, LYS2, met15Δ, leu2Δ::LEU2-*$P_{MYO2}$*-rtTA-Adh1t, ho::HIS5-*$P_{TETO4}$*-I-SceI-Cyc1t, ura3Δ::URA3-*SSAreporter(control, no cut-site, with mCherry-ssCLN2PEST) |
| yTY147a ('SSA+ RFPdegron' in figures) | SSA (RFP degron) | *MATα, his3Δ, LYS2, met15Δ, leu2Δ::LEU2-*$P_{MYO2}$*-rtTA-Adh1t, ho::HIS5-*$P_{TETO4}$*-I-SceI-Cyc1t, ura3Δ::URA3-*SSAreporter(perfect homology, with mCherry-ssCLN2PEST) |
| yTY149b ('SSA *dnl4Δ*' in figures) | SSA (perfect homology), *dnl4Δ* | *MATα, his3Δ, LYS2, met15Δ, leu2Δ::LEU2-*$P_{MYO2}$*-rtTA-Adh1t, ho::HIS5-*$P_{TETO4}$*-I-SceI-Cyc1t, ura3Δ::URA3-*SSAreporter(perfect homology), *dnl4Δ::KANMX4* |

**Table S1. Yeast strain descriptions and genotypes.**

| Strain Description | Age group at the start of dox treatment | Replicate | Number of cells used in repair efficiency calculation | Number of unrepaired cells at the start of dox treatment | Number of repaired cells at the start of dox treatment |
|---|---|---|---|---|---|
| SSA | old | 1 | 21 | 32 | 4 |
| SSA | old | 2 | 18 | 29 | 6 |
| SSA | young | 1 | 33 | 39 | 1 |
| SSA | young | 2 | 31 | 31 | 3 |
| SSAcontrol | old | 1 | 30 | 41 | 0 |
| SSAcontrol | old | 2 | 31 | 38 | 0 |
| SSAcontrol | young | 1 | 21 | 22 | 0 |
| SSAcontrol | young | 2 | 44 | 44 | 0 |
| SSAheterology | old | 1 | 20 | 42 | 0 |
| SSAheterology | old | 2 | 32 | 43 | 2 |
| SSAheterology | old | 3 | 21 | 33 | 1 |
| SSAheterology | old | 4 | 20 | 31 | 2 |
| SSAheterology | old | 5 | 16 | 28 | 0 |
| SSAheterology | young | 1 | 46 | 48 | 0 |
| SSAheterology | young | 2 | 40 | 42 | 0 |
| SSA *dnl4*Δ | old | 1 | 38 | 44 | 5 |
| SSA *dnl4*Δ | old | 2 | 22 | 30 | 2 |
| SSA *dnl4*Δ | young | 1 | 25 | 25 | 3 |
| SSA *dnl4*Δ | young | 2 | 27 | 27 | 3 |
| SSAcontrol+ RFPdegron | old | 1 | 32 | 40 | 0 |
| SSAcontrol+ RFPdegron | old | 2 | 26 | 32 | 0 |
| SSA+ RFPdegron | old | 1 | 32 | 44 | 6 |
| SSA+ RFPdegron | old | 2 | 23 | 41 | 5 |
| SSA+ RFPdegron | old | 3 | 26 | 36 | 5 |

**Table S2. Strains, age groups, and experimental replicates/cells used for SSA repair efficiency calculations.** The last column refers to the cells that, at the beginning of dox treatment, were both unrepaired (based on YFP) and in the appropriate age range. For a cell to be included in the repair efficiency calculation, it had to not only satisfy these conditions, but also be alive 5 hours after removal of doxycycline.

| Strain Description | Age group at the start of dox treatment | Replicate | Number of cells used in repair efficiency calculation | Number of unrepaired cells at the start of dox treatment | Number of repaired cells at the start of dox treatment |
|---|---|---|---|---|---|
| SSA+ RFPdegron | young | 1 | 14 | 14 | 1 |
| SSA+ RFPdegron | young | 2 | 23 | 24 | 3 |

**Table S2 (continued). Strains, age groups, and experimental replicates/cells used for SSA repair efficiency calculations.**
